# Trajectory matching-random walk feasibility sampling for simulating heterogeneous dynamics in mechanistic models

**DOI:** 10.1101/2025.09.26.678800

**Authors:** Fatemeh Beigmohammadi, Jordan J.A. Weaver, Solène Hegarty-Cremer, Cailan Jeynes-Smith, Amber M. Smith, Morgan Craig

**Affiliations:** Sainte-Justine University Hospital Research Centre, Montréal, Québec, Canada; Department of Mathematics and Statistics, Université de Montréal, Montréal, Québec, Canada; Department of Pediatrics, University of Tennessee Health Science Center, Memphis, TN USA

**Author notes:** Corresponding author: Morgan Craig.

**Keywords:** virtual patient cohorts, trajectory matching, random walk feasibility sampling, plausible populations, mechanistic mathematical models

## Abstract

The inherent heterogeneity of complex biological systems makes it difficult to experimentally and clinically explore individual outcomes. Mechanistic mathematical models are essential tools for studying such heterogeneity. Thus, there is increasing interest in integrating newer mechanistic model-based techniques, like virtual patient cohorts and virtual clinical trials, within experimental and regulatory pipelines to probe relationships that may be difficult or impossible to ascertain through traditional wet-lab or clinical experimentation alone. Approximate Bayesian Computation (ABC) is an attractive method for generating virtual patient cohorts and running virtual clinical trials. However, ABC-based approaches can become computationally demanding when stringent acceptance criteria are imposed. In response, we developed a model-based technique called trajectory-matching random-walk feasibility sampling (TM-RWFS) that captures the variability of complex biological systems by constraining model trajectories between the upper and lower bounds of available data to generate heterogeneity. By testing the method’s performance on existing mechanistic models, we show that TM-RWS generates parameter sets whose simulated trajectories reproduce the observed variability in biological systems of varying complexity while maintaining computational efficiency. Thus, TM-RWFS is a fast, new approach for generating heterogeneity in mechanistic mathematical models with implications for model-based experimental design, virtual patient cohorts, and virtual clinical trials.

## 1. Introduction

Mathematical models of disease progression and treatment have become indispensable tools for understanding biological systems and informing clinical decision-making. One challenge of modelling disease progression and treatment beyond available data lies in capturing the diversity of biological trajectories^1–7^, which arises as a consequence of numerous factors, including age, sex, comorbidities, and immune status. Characterizing this variability experimentally or clinically is often constrained by time, cost, and ethical considerations, making it generally infeasible to explore the full range of plausible scenarios through data collection alone. Mechanistic mathematical models offer a powerful means of describing biological heterogeneity and are therefore essential tools to probing variability that would be impractical to assess experimentally or clinically^8–13^.

Several approaches quantify optimal parameter sets and capture heterogeneity in biological data through fitting mechanistic model outputs using global optimization algorithms, such as adaptive simulated annealing, genetic algorithms, or sampling algorithms such as Metropolis-Hastings^14–16^. These methods can be combined with bootstrapping, while Bayesian approaches such as like Approximate Bayesian Computation (ABC) or Markov Chain Monte Carlo (MCMC) can be used to estimate posterior parameter distributions^17–23^. Across these approaches, model parameters are repeatedly evaluated to identify parameter sets that fit the data, thereby characterizing population-level heterogeneity. However, an important goal of in silico clinical trial simulation is to identify plausible model trajectories that extend beyond available data or perform well under conditions of limited data, enabling the use of virtual patients to predict outcomes. Prior studies have used simulated annealing^7^, ABC^10^, and ABC-MCMC^24,25^ methods to ensure model trajectories remain between upper and lower data bounds. ABC and ABC-MCMC avoid direct likelihood evaluation but can increase in execution time when stringent acceptance criteria substantially increase the computational effort required to obtain acceptable solutions^17–19,24,26–31^. Despite efforts to improve computational efficiency^25,32^, such as sequential Monte Carlo ABC (ABC-SMC) and parallel candidate proposals^18,19,25,31,33–35^, these methods rely on approximations (i.e., summary statistics) and assumptions (i.e., prior choice, distance metrics, and tolerance thresholds) that can influence the resulting inference^33^, particularly when data are sparse and lack fine granularity. In response, we developed a trajectory-matching-random walk feasibility sampling algorithm that directly leverages the observed data by constraining model simulations to lie within their upper and lower bounds, improving both biologically plausibility and computationally efficiency. Here, heterogeneity refers to variability in parameter values within the feasible population, which results in variability in model trajectories.

Our method simulates model outputs and compares them to experimental data using parameters drawn from normal distributions whose means are updated to the most recently accepted values, with standard deviations determined from previously estimated best-fit values. Sample acceptance occurs when model-simulated trajectories lie within the upper and lower bounds of the observed experimental data, a criterion we term trajectory matching (TM). We consider several previously calibrated biomedical ODE models^3,5–7,30,36–42^ and define the parameter sets that represent individual trajectories. By accepting parameter sets whose trajectories satisfy the specified bounds, a range of individualized model outcomes are generated (e.g., virtual patients) that reflect the variability represented by the population data. Using these examples, we evaluate acceptance rates, computational performance, and the ability of TM-RWFS to generate feasible parameter populations that capture complex, dynamic behaviours in disease models. Importantly, TM-RWFS displayed fast execution across all studied examples, supporting its utility for rapid virtual patient generation. Together, these features support the use of mechanistic models for in silico experimentation, including modelling disease outcomes with virtual patient cohorts and informing effective treatments through virtual clinical trials^10,43–47^.

## 2. Methods

### 2.1. TM-RWFS pseudocode

A flow chart of the TM-RWFS algorithm is given in **Figure 1**. For each of the *M* runs, the proposal means are initialized using specified initial parameter values. To initialize the algorithm, let 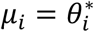 for the first iteration, where *μ*_*i*_ is normal distribution mean of *i*^th^ parameter (i.e., when *n* = 1, 1 ≤ *n* ≤ *N*, with *N* the total number of iterations). 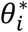 is then updated at each subsequent iteration based on newly accepted parameter sets as follows.

**Figure 1.**
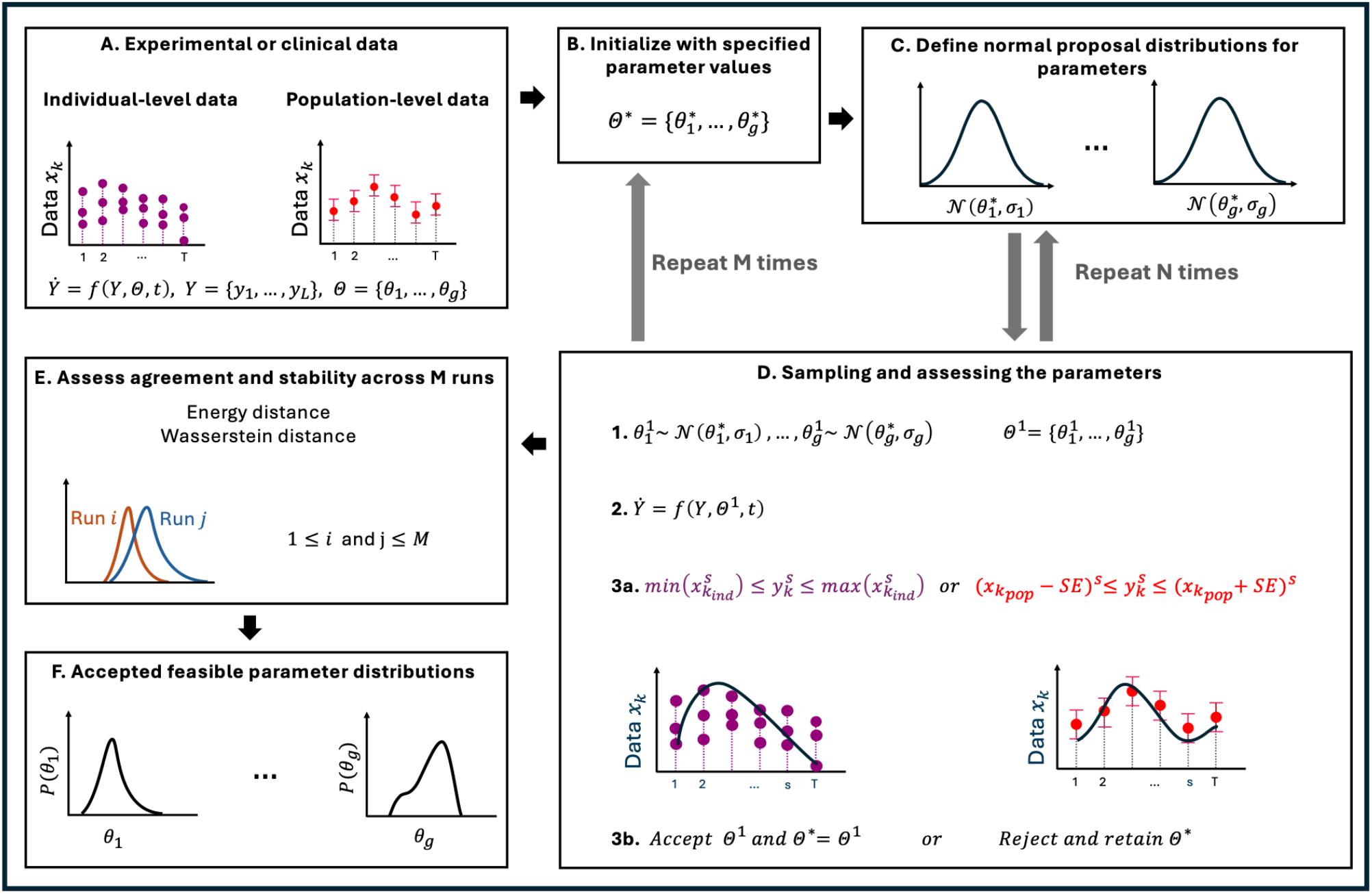
Overview of trajectory-matching random-walk feasibility sampling algorithm. A) The TM-RWFS algorithm starts with experimental or clinical data x_k_, corresponding mathematical model Y, and previously estimated parameters 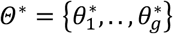. B and C) To initialize the algorithm, define the means of normal proposal distributions as 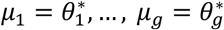. Then, repeat steps 1 to 3 for N iterations to generate 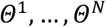. D) Update each mean in normal proposal distributions if step 3a is fulfilled for a (pre-determined and) sufficient number of time points. Repeat the procedure for M pre-specified runs using the specified initial parameter values. E) Assess agreement and stability across runs by evaluating the distance between proposal distributions. F) Construct accepted feasible parameter distributions using parameter sets associated with accepted trajectories.

***Step 1***. For 2 ≤ *n* ≤ *N* and 1 ≤ *m* ≤ *M*, where *M* is total number of parallel runs, sample candidate parameters from normal proposal distributions:

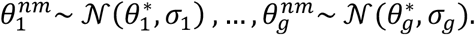

Here, {*σ*_1_, *σ*_2_, …, *σ*_*g*_} is the set of standard deviations of the first parameter (index 1) to the last parameter (index *g*), respectively, that determines the width of the normal distribution *N* and 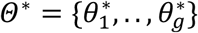 is the vector of initial parameters values.

***Step 2***. Simulate *Y* from the model ℳ given candidate parameters 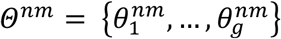.

***Step 3a***. Assess the feasibility of predicted outputs *Y* with respect to the data at each time point *s*. This is performed by checking whether the predicted model trajectory falls within the bounds defined in Eq. (8) or Eq. (9). If the new parameter set meets these acceptance criteria, i.e., Eqs. (8) or (9) satisfied at the pre-defined number *q* of the *T* selected time points, where 1 ≤ *q* ≤ *T*, and if all additional case-specific feasibility constraints are satisfied, accept *Y* and the candidate parameter set *Θ*^*nm*^ and update normal distribution for the next iteration to be

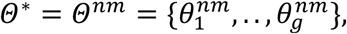

***Step 3b***. Otherwise, reject the candidate parameter set and continue with the previously accepted parameter values in normal distributions:

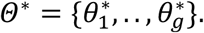

Note that no likelihood-based Metropolis-Hastings ratio is evaluated; acceptance or rejection is determined solely by the specified feasibility criteria.

Repeat **Steps 1**-**3** for *N* iterations in each of the *M* runs.

***Step 4***. Assess agreement and stability across the pre-specified *M* runs using acceptance rates and the diagnostic measures. Energy distance is used to quantify agreement between the joint accepted parameter populations from two TM-RWFS runs.

Let *φ* and *ψ* define d-dimensional random parameter vectors associated with runs *i* and *j*, respectively, and let *φ*′ and *ψ*′ denotes independent copies of *φ* and *ψ*. The energy distance is defined as:

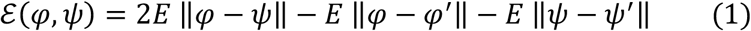

where *E*[·] denotes the average over all possible realizations and ‖ · ‖ denotes the Euclidean norm. In TM-RWFS, each realization of *φ* and *ψ* corresponds to a complete accepted parameter vector from the corresponding run. Energy distance compares the joint parameter populations to check if they come from same distributions. Thus, the run with the lowest mean energy distance is selected as the representative run from the *M* total runs. We then construct the accepted feasible parameter distributions {*P*(*θ*_1_), *P*(*θ*_2_), …, *P*(*θ*_*g*_)} from the parameter sets of the selected representative run; the resulting empirical parameter distributions describe the accepted feasible population.

### 2.2. Computational implementation

All simulations and analyses were performed in Julia (version 1.10). The implementation used several open-source packages, including *DelayDiffEq* and *Sundials* for solving differential equations; *Distributions, StatsBase*, and *KernelDensity* for statistical routines; *PlotlyJS* for interactive visualizations; and *DataFrames* for data management. Additional packages, including *Interpolations*, Q*uadGK*, and *Parameters* were used for interpolation, numerical integration, and parameter handling.

### 2.3. Case Study 1: Lotka Volterra model

The Lotka-Volterra (LV) model is a deterministic system commonly used in ecology to describes the interaction between prey (*x*) and predator (*y*) populations. Here, we used the LV model given by:

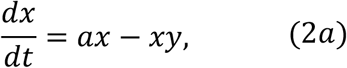

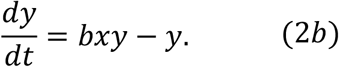

Parameters *a* and *b* describe the growth rate of the prey, *x*, in the absence of predators, *y*, and the rate of predator reproduction due to interaction with prey, respectively ^31,48,49^. Toni et al. applied ABC and its variants to the LV model to compare their efficiency based on acceptance rates. For this purpose, they generated synthetic data for both species, *x* and *y* (*D*_*x*_ and *D*_*y*_), by adding Gaussian noise, (N(0, 0·5^2^)) to simulated trajectories at eight randomly selected time points.

Because Toni et al.^31^ reported similar performance between ABC and ABC-MCMC, we limited our benchmarking comparisons to rejection ABC and TM-RWFS. For this, we implemented the rejection ABC procedure following the approach described by Toni et al.

When a system exhibits well-defined extrema (e.g., stable oscillations), synthetic data bounds can be constructed by applying a fixed percentage of uncertainty (e.g., 20%) around those points. Thus, to compare TM-RWFS to ABC, we selected synthetic data points based on the extrema of the model’s prediction for one of the components (here, *x*) to generate trajectories and parameter distributions. Using the parameters as in Toni et al., the component *x* has two minima and three maxima over fifteen days. Therefore, we defined bounds at five time points according to:

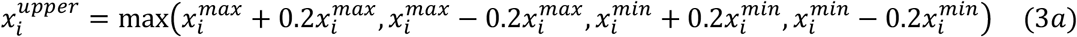

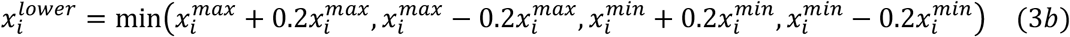

Let the set of these points be *D*_*extrema*_. Parameters *a* and *b* were then sampled from normal distributions with means *μ*_*a*_ = *μ*_*b*_ = 1 and variances *σ*_*a*_^2^ = *σ*_*b*_^2^ = 0·1. As rejection ABC and TM-RWFS use different sampling structures and acceptance procedures, an identical comparative implementation is not possible. We therefore compared the methods using the common criterion available to both approaches, namely the discrepancy between simulated trajectories and synthetic data. Thus, we used the same sum-of-squared-errors distance function and acceptance threshold of *ϵ* = 4·3 for both methods to evaluate the performance of each algorithm under this distance-based criterion; we note that our results should thus not be interpreted as demonstrating the intrinsic superiority of one algorithm over the other (Table S1). In this example, the model trajectory fell within the constructed bounds at only three of the eight time points. Therefore, a strict comparison using the same eight noisy observations is not generally feasible without altering the TM-RWFS acceptance criterion, because the bounds may not define a jointly feasible trajectory. We instead compared the methods using the common distance function and acceptance threshold available to both approaches (see “Scaling the observed data” and Figures S10A and S11 in the Supplementary Information for comparisons using identical data points and observation times).

While extrema are often informative points in a model’s dynamics, other dynamical features, including waning rates and plateaus, may be more salient properties for a given system. In such cases, time points must be selected strategically. See the Supplementary Information for a discussion of some strategies for generating the synthetic data and their influence on TM-RWFS performance, including how varying the level of agreement between data and model affects the resulting trajectories and parameter distributions (Table S8, Figure S12, S13; the importance of time point selection is further discussed in Section S6 of Supplementary information (Table S5 and Figure S7) where we evaluated how different choices of trajectory-matching time points affected the resulting accepted parameter sets and trajectories using the first influenza-infection model).

### 2.4. Case Study 2: Models of influenza virus infection

The second model to which we applied TM-RWFS is a system of ordinary differential equations (ODEs) describing the viral kinetics of influenza A infection^50,51^. The model includes four variables: susceptible epithelial cells (*T*), eclipse infected cells (*I*_1_), productively infected cells (*I*_2_), and viral load (*V*):

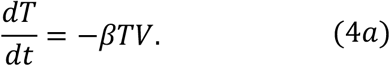

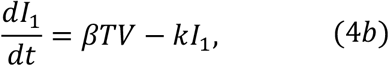

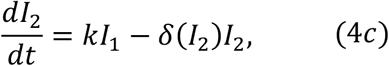

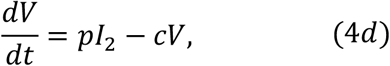

where 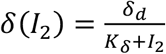, *δ*_*d*_ represents the maximum rate of infected cell clearance, and *K*_*δ*_ denotes the half-saturation constant. The subscript *d* indicates that the clearance process is density dependent.

Parameters were fit to viral loads measured from the lungs of infected groups of mice over the course of 12 days using adaptive simulated annealing (ASA). Smith et al.^52^ reported the best fit values and 95% CIs (confidence intervals) of each parameter (Table S2). The value of *k*, the rate of transition through the eclipse phase, must be restricted between 4 and 6 to ensure biological relevance. This model was then extended by Myers et al.^36^ to study CD8^+^ T cell immune responses by adding equations for effector (*E*) and memory (*E*_*M*_) CD8^+^ T cells:

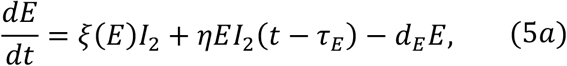

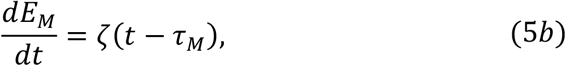

where 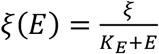 and *η* describes the infiltration and expansion of effector CD8^+^ T cells, respectively, and *ζ* describes the generation of memory CD8^+^ T cells. Eq. (4*c*) is also changed to reflect the clearance of infected cells by T cells:

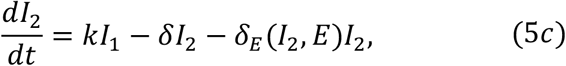

with 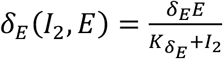. The extended version has 14 parameters. Myers et al.^36^ similarly obtained best-fit parameter values and their 95% CIs by fitting their model using simulated annealing and bootstrapping viral load and total CD8^+^ T cell count data from days 0 to 12 (*T* = 13). Most parameter values were deemed to be well defined, while a select few were unidentifiable (i.e., the accepted distribution was uniform)^36^ (Table S3).

### 2.5. Case Study 3: Model of tumour growth after combination immunotherapy

Finally, we applied TM-RWFS to a cancer immunotherapy model describing the tumor response to a combination of an oncolytic virus and injections of dendritic cells (DCs)^53^. The treatment schedule considered includes injections of 2·5 × 10^9^ oncolytic virions on days 0, 2, and 4, and 10^6^ dendritic cells on days 1, 3, and 5^51^. The model tracks uninfected tumor cells (*U*), tumor cells infected by an injected oncolytic virus (*I*), oncolytic virus (*V*), naïve CD8^+^ T cells (*A*), tumor antigen-specific CD8^+^ T cells (*T*), and injected dendritic cells (*D*):

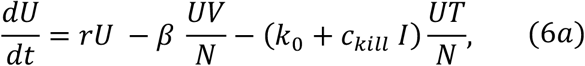

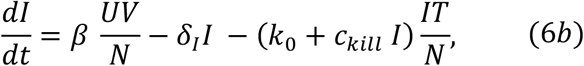

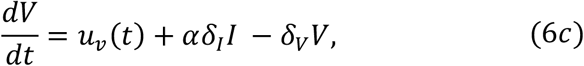

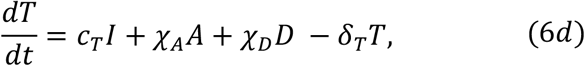

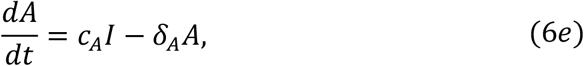

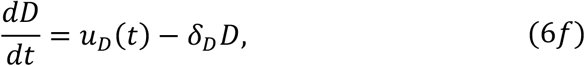

This model has fourteen parameters, six of which (the net tumor growth rate, *r*, infection rate *β*, naïve T cell recruitment rate *c*_*A*_, T cell recruitment rate *c*_*T*_, enhanced T cell cytotoxicity rate, *c*_*kill*_, and the rate of T cell simulation by DCs *χ*_*D*_) were estimated and reported with corresponding 95% credible intervals (CrI; Table S4). Confidence intervals and credible intervals have different interpretations. In the frequentist framework, a 95% *CI* is defined such that 95% of the calculated intervals would contain the true parameter value if the experiment were repeated many times. However, no probability can be assigned to any specific interval. In contrast, a 95% *CrI* in the Bayesian framework indicates that there is a 95% probability that the true parameter value lies within the interval, given the observed data. In Barish et al.^40^, parameters were estimated hierarchically using simulated annealing based on aggregated data. Here, we generated the necessary distributions about the best-fit values of the six parameters with CrIs in a single step using TM-RWFS, i.e., without employing the hierarchical fitting procedure.

## 3. Results

### 3.1. TM-RWFS algorithm

To generate plausible parameter sets and model trajectories within data-derived constraints, we developed trajectory-matching RWFS. TM-RWFS is a model-based algorithm combines random-walk parameter proposals with a trajectory-matching acceptance criterion. Experimental data and a mathematical model form the backbones of TM-RWFS. To apply the method, we suppose the availability of (1) observed data for at least one model component, (2) point estimates of parameters, and (3) confidence intervals (CIs). However, scenarios may arise where one or more of these elements are missing, e.g., when CIs are not reported, parameter estimates have low confidence, or estimating plausible bounds on the data is not possible. In such cases, TM-RWFS can still be applied, but its sampling efficiency and exploration of the feasible parameter region may be affected (see Supplementary Tables S6 and S7).

The TM-RWFS procedure is as follows. Assume that we have an existing mechanistic model describing a biological system with previously defined biologically plausible bounds on its relevant parameters. Define this mechanistic model, ℳ, as the system of differential equations:

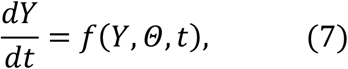

where *Y*(*t*) is the set of *L* model state variables *Y*(*t*) = {*y*_1_, · ·, *y*_*L*_}, *Θ* is the set of *g* model parameters *Θ* = {*θ*_1_, · ·, *θ*_*g*_}, and *t* is time. Simulation of ℳ yields time-dependent trajectories, *y*_*i*_(*t*), that predict the model behavior with parameters *Θ*.

Let *x*_*k*_ 1 ≤ *k* ≤ *L* be the set of experimentally observed outcomes and let *T* denote the total number of measurements. The first step of TM-RWFS is to define the response interval given the upper and lower bounds for each *x*_*k*_. If there are individual-level data available 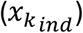, we define the response interval to be

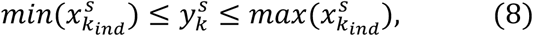

where 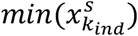 and 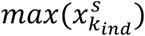 are the minimum and maximum values, respectively, of each data point at the discrete time point *s*, for 1 ≤ *s* ≤ *T*. Alternatively, if the observed data are reported as summary statistics for the population, 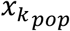 represents the population-level data, and the response interval is instead defined as

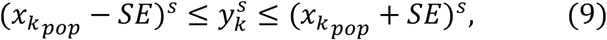

i.e., the *mean* ± *standard error* (*SE*). TM-RWFS uses these response intervals as acceptance criteria.

Once response intervals are established, several parameters must be specified. The first is the number of iterations, *N*. The second is the pre-specified number of parallel runs *M*. The appropriate values of *N* and *M* depend on a given model and computational setting, but they are generally in the range of [10^4^, 10^6^] and [2,5], respectively. We performed *M*parallel runs to assess agreement, stability, and consistency of the accepted feasible parameter populations across repeated TM-RWFS runs.

For the random-walk sampling step, normal proposal distributions around parameter means must be defined. These proposal distributions are assumed to be normal, N(*μ, σ*^2^). We set the distribution means to be the best-fit parameter estimates, 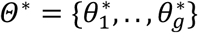. These best-fit values are the set of parameters that best minimize the difference between model predictions and the observed data (see Discussion) and are assumed to be available before running TM-RWFS.

Thus, the resulting normal proposal distribution of parameter *θ*_*i*_ is given by:

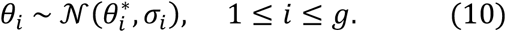

where 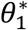 is the best fit value of parameter *θ*_*i*_, and 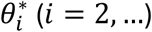 is the current proposal mean, initialized using the specified initial value and updated to the most recently accepted value.

Let 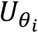 and 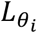 denote the upper and lower bounds, respectively, of the estimates on parameter *θ*_*i*_, which are derived from available confidence intervals. The standard deviation, *σ*_*i*_, is defined according to the difference between these upper and lower bounds:.

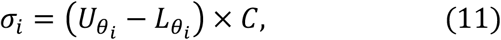

where *C* ∈ [0·05,0·1] is a user-specified proposal-scaling coefficient that influences acceptance and exploration of parameter space. If 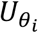 and 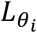 ∈ [10^−2^, 10^2^), sampling is performed on a linear space. Otherwise, sampling is performed in logarithmic space, and the logarithmic values of 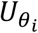 and 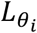 is used to calculate *σ*_*i*_. Typical values of *σ*_*i*_ fall between [0·05, 0·5]. When sampling in logarithmic space produces a very small *σ*_*i*_, we neglect the coefficient *C* and simply define *σ*_*i*_ to be 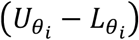. When confidence intervals (CIs) are not available, the standard deviation can be defined as a constant value *σ*_*i*_ = 0·1 across all normal distributions (see Supplementary Information).

The proposal distributions in Eq. (10) guide the exploration of the parameter space, affecting how candidates are generated, whereas parameter bounds, trajectory-matching criteria, and additional constraints define the feasible region. Parameter sets whose simulated trajectories satisfy the specified response intervals and any additional case-specific constraints are retained to construct the feasible set of parameter sets.

TM-RWFS follows a random-walk sampling strategy in which candidate parameters depend on the most recently accepted values. A candidate parameter set is accepted when its model-predicted trajectory lies within the specified upper and lower data bounds at the required measurement times. Otherwise, the candidate is rejected, and the previous proposal mean is retained.

### 3.2. Assessing the agreement and stability of TM-RWFS

As each TM-RWFS run is subject to random parameter proposals, some run-to-run variability is expected. Thus, after *N* iterations, we used energy distance^55^ Eq. (1) to assess the agreement and stability of the accepted feasible parameter populations across the pre-specified *M* runs. Energy distance quantifies the agreement between the joint multivariate parameter populations. Wasserstein distance^54^, another metric that quantifies agreement between the marginal parameter distributions across runs, gave essentially the same results (See SI, section S12). For generating virtual populations, a single representative was selected to be the run with the smallest mean energy distance to the remaining runs, representing the run whose joint parameter population was most similar to the other runs, using a pre-specified quantitative criterion.

### 3.3 Testing the performance of TM-RWFS for complex biological systems

To evaluate the performance of TM-RWFS, we selected four models of varying complexity and precision in parameter estimates (see Tables S1-S4). For all tested models (except the one in Case Study 1), the original studies obtained parameter distributions using simulated annealing-based algorithms. We systematically evaluated the performance of the TM-RWFS method by beginning with the model that had the fewest equations and parameters and then progressing to more complex models. In each case, we verified we had correctly implemented the model by reproducing figures in the original papers^36,40,52^ before applying TM-RWFS.

#### Case Study 1: Performance of TM-RWFS compared with ABC methods on the Lotka Volterra Model

We began by benchmarking the TM-RWFS algorithm against the standard ABC rejection algorithm applied to the Lotka–Volterra (LV) model^31,48,49^. We first compared TM-RWFS with ABC with the goal of quantifying the computational cost of generating approximately 1000 accepted samples from each of the algorithms. We compared the resulting parameter distributions of *a* and *b*, and the corresponding trajectories of *x*. The same comparison could be made using trajectories of *y*; however, since the process is identical, we arbitrarily chose to focus on *x*. Owing to the simplicity of the Lotka-Volterra model and the sampling of only two parameters, repeated TM-RWFS runs produced highly similar accepted feasible parameter populations. We therefore used a single run (*M* = 1) for this case study and did not apply the representative-run selection procedure.

To account for differences in sampling structure and acceptance mechanisms, we compared TM-RWFS and rejection ABC by comparing the differences between simulated trajectories and synthetic data in each method, given the same sum-of-squared-errors distance function and acceptance threshold, *ϵ*. Under the specified comparison conditions, TM-RWFS required fewer simulations to generate approximately 1,000 accepted samples, terminating in 9 seconds compared with 7 hours for rejection ABC (**Table 1**). The resulting range of *ϵ* values calculated for TM-RWFS was [3·8, 5·5]. To enable comparing with ABC, we focused on the subset of accepted parameters with *ϵ* ≤ 4·3, as in Toni et al. However, the remaining parameter sets (with *ϵ* values in the range *ϵ* ≤ 5·5) produced broader feasible parameter distributions, leading to greater heterogeneity in the resulting trajectories, even though their *ϵ* values were only slightly above the comparison threshold (**Figure 2**).

**Table 1.** Comparison of ABC and TM-RWFS performance on the Lotka-Volterra model. N indicates the total number of simulations, and AR is the acceptance rate. Both methods used the same distance function (sum of squared errors, SSE) and threshold (ϵ = 4·3). These results describe computational performance under the specified acceptance criteria and should not be interpreted as demonstrating intrinsic computational superiority of TM-RWFS over ABC.

| Algorithm | N | Data type | # Data points | Distance function | Execution time | # Accepted | $\epsilon$ | AR (%) |
| --- | --- | --- | --- | --- | --- | --- | --- | --- |
| ABC | $1.4 \times 10^7$ | $D_x, D_y$ | 8 | SSE | 7 hours | 912 | 4.3 | 0.006 |
| ABC-SMC | 12866 | $D_y$ | 8 | SAE | 75 seconds | 1000 | 4.3 | 7.7 |
| TM-RWFS | $10^4$ | $D_{\text{Extrema}}$ | 5 | SSE | 9 seconds | 1263 | 4.3 | 12 |

**Figure 2.**
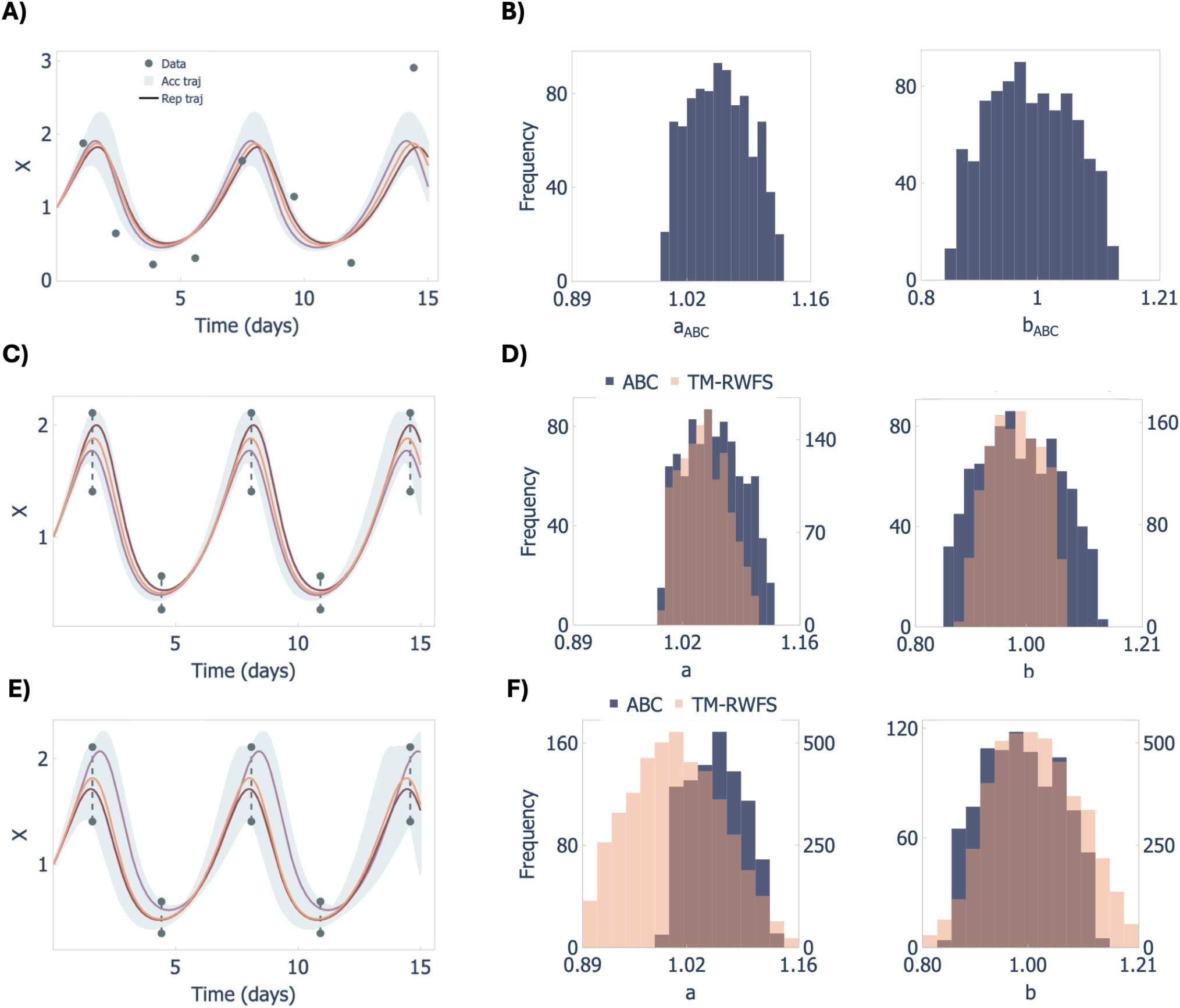
Benchmarking results comparing ABC and TM-RWFS for the Lotka-Volterra model. A) Accepted model trajectories for component x using the synthetic data D_x_, B) Accepted parameter distributions for parameter a and b, obtained using the ABC algorithm with an acceptance threshold of ϵ ≤ 4·3. C-D) Same as panels A-B but using the TM-RWFS algorithm with an acceptance threshold of ϵ ≤ 4·3 and D_Extrema_. E-F) Same as panels A-B but with a relaxed acceptance threshold (ϵ ≤ 5·5), illustrating how the TM-RWFS recovers broader and more diverse trajectories while maintaining good agreement with data.

We also tested ABC-SMC on the LV model using the pyABC package, i.e., a Python implementation of Approximate Bayesian Computation Sequential Monte Carlo (ABC-SMC)^19,31,35,56,57^, with an execution time of 75 seconds and acceptance rate of 7.7% (**Table 1**). Moreover, although ABC-SMC outperforms ABC, the accuracy of the inferred parameter distributions and the extent to which the resulting trajectories are biologically meaningful depend strongly on several factors, including the choice of prior distributions, the definition of the distance function, the threshold (*ϵ*) range, and the amount of data incorporated into the algorithm (see Section S10 in the Supplementary Information). In some cases, ABC-SMC outperformed TM-RWFS, especially when prior biological knowledge can be used to constrain plausible parameter ranges and the model is sufficiently robust to match the data well, enabling the generation of parameter sets that closely reproduce the observed dynamics (see Section S9 and S10 in the Supplementary Information). The acceptance-related metric (*ϵ*) is reported here solely to enable comparison with existing ABC variants. In TM-RWFS, computational performance is additionally assessed using execution time and the number of simulations required to generate accepted feasible parameter sets.

#### Case Study 2: Model of influenza virus infection and CD8^+^ T cell immune responses

Comparing TM-RWFS to ABC variants using a straightforward model like the Lotka-Volterra model allowed us to validate the computational performance of our algorithm under the specified comparison conditions. We now extend our approach to more mechanistic and biologically realistic models to assess its performance in increasingly complex systems. Specifically, we applied TM-RWFS to a model of influenza virus infection that includes density-dependent clearance of infected cells^52^. To perform TM-RWFS, we used *M* = 5 parallel runs, each with *N* = 2 × 10^5^ iterations. At each iteration, a candidate parameter set *Θ* = {*β, p, c, k, δ*_*d*_, *K*_*δ*_} was sampled from normal proposal distributions centred on the most recently accepted parameter values (see Methods). For a candidate solution to be accepted, predicted viral loads were required to be within the data’s upper and lower bounds (*Eq*· (7)) with *q* = 11 for the *T* = 11 time points. Accepted model trajectories represented 1184 of the total simulations (*N* = 2 × 10^5^), yielding an acceptance rate of 0·6 %. Resulting trajectories and illustrative distributions are shown in **Figure 3**. TM-RWFS completed in 13 minutes and produced parameter distributions consistent with Smith et al., demonstrating both significant computational efficiency and robustness. The accepted parameter distributions closely reproduced the heterogeneity reported in Smith et al^22^ (Figure S1), which was further confirmed by recreating scatter plots between parameter pairs (Figure S2). We also implemented this model in pyABC package. Given comparable scenarios, TM-RWFS and pyABC showed comparable computational performance, though the acceptance threshold *ϵ* and prior definitions have a considerable effect on execution time and on the posterior distributions of some model parameters (see Section S10 in the Supplementary Information). In this model, *M* = 5 was selected as representative run with mean energy distance 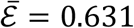.

**Figure 3.**
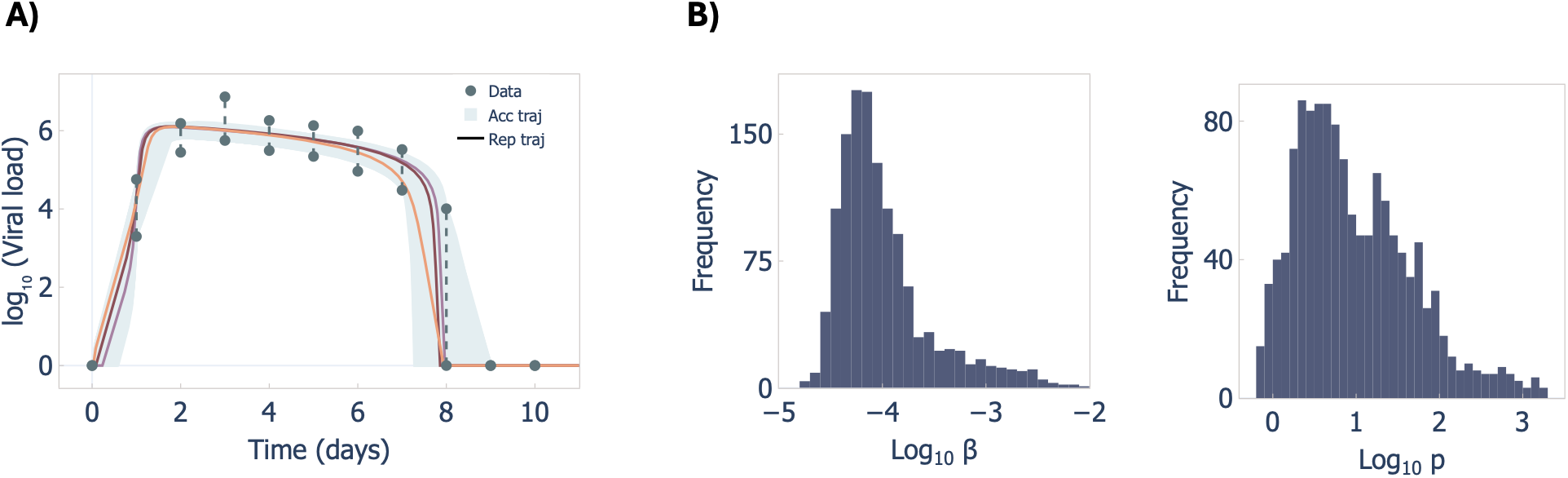
Outputs of the TM-RWFS algorithm on the density dependent viral kinetic model. A) Accepted trajectories for virus dynamics. B-left) Accepted feasible parameter distribution of virus infection rate β, and B-right) virus production rate p.

We then applied TM-RWFS to the extended model of murine influenza infection describing viral titers and CD8+ effector and memory T cell concentrations^36^ (see Methods). TM-RWFS was implemented by performing *M* = 5 parallel runs, each with *N* = 15 × 10^4^ iterations. At each iteration, a candidate parameter set 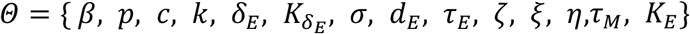 was sampled from normal proposal distributions. Because the data were a time-series with 10 independent individuals at each collection time, a candidate paameter set *Θ* was accepted only if the simulated outcomes for the model solutions for the virus fell within the virus data bounds and for the *CD*8^+^ *T* cell fell within the *CD*8^+^ *T* cell data bounds, for *q* = 13 total timepoints. TM-RWFS yielded an acceptance rate of 0·9 % (see resulting accepted trajectories in **Figure 4A-4B** with illustrative distributions in **Figure 4C**; full parameter distributions are in Figures S3-S6) and completed in 10 minutes. Together, these results show that even with increased model complexity, TM-RWFS maintains a high degree of accuracy while maintaining computational efficiency, indicating its robustness and adaptability to more complex systems. The higher acceptance rate observed for this model, relative to the previous influenza model, was associated with the choice of the proposal scaling coefficient *C*, which was set to 0·07 instead of 0·1. (see Discussion). Distributions of the rest of the parameters are shown in Figures S3-S6. For this case study, *M* = 2 was selected as representative run with mean energy distance 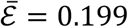.

**Figure 4.**
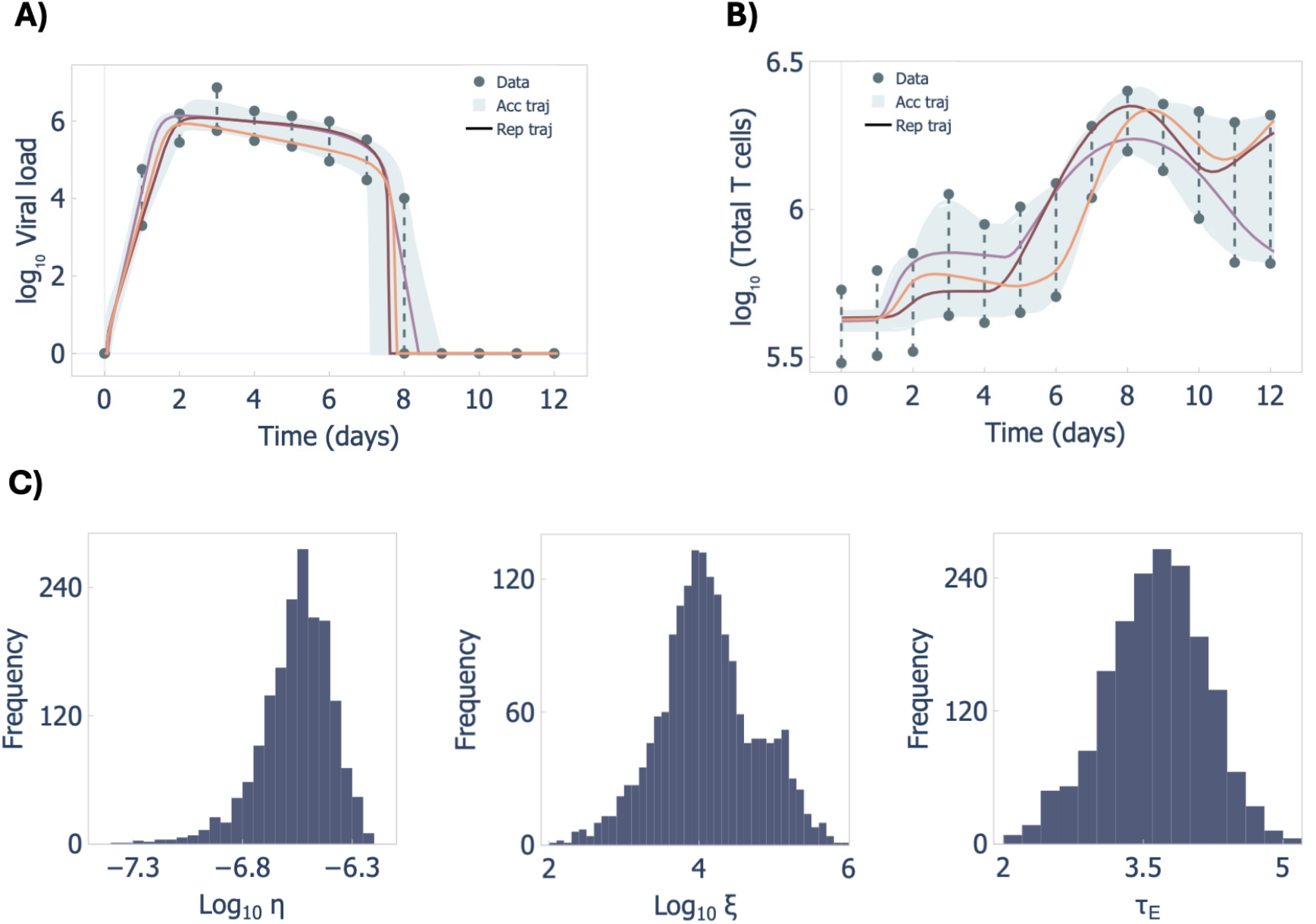
Output of TM-RWFS algorithm on extended model of inffuenza infection including CD8+ T cell dynamics. A) Accepted trajectories for viral dynamics and B) total number of CD8^+^ T cells. C-left) Accepted feasible parameter distribution of the CD8_E_ expansion rate η, C-middle) CD8_E_ infiltration rate ξ and C-right) delay in CD8_E_ expansion rate τ_E_·

#### Case Study 3: Model of cancer immunotherapy

Lastly, we applied TM-RWFS to a model describing a combination cancer immunotherapy model^40,53,58^. To implement the TM-RWFS algorithm, we performed *M* = 5 parallel runs, each with *N* = 12 × 10^4^ iterations. At each iteration, a candidate parameter set *Θ* = {*r, β, χ*_*D*_, *c*_*A*_, *c*_*T*_, *c*_*kill*_} was sampled from normal proposal distributions. In this case, only population-level data were available. Thus, to ensure acceptance (*Eq*· (8)), simulated trajectories of the candidate parameter set had to remain within the tumor volume bounds defined by *mean* ± *SE* (*standard error*), with a 10% tolerance added to the standard error as a case specific feasibility choice. In the original paper, the model tracks well with the experimental data after day 7 but fails to adequately capture the earlier dynamics from days 0-7. Accordingly, we restricted the acceptance window to *q* = 22 time points and excluded the first seven points to avoid constraining parameter acceptance during a period that the model was not designed to accurately reproduce. We further imposed a feasibility constraint so that predicted viral loads remained below 1 × 10^9^ virions at day 30, consistent with the viral dynamics reported in Wares et al.^53^. This constraint effectively incorporates additional biological knowledge of viral dynamics into the selection of accepted trajectories, as in the previous Case Study (influenza infection) where we included information about both viral load and T cell dynamics. Given that we only required that model trajectories match 22 of 30 time points, the value of *N* was 12 × 10^4^. This also reflects the lower complexity of the mathematical model. Accepted model trajectories (**Figure 5A**; illustrative distributions in **Figure 5B**) comprised 2% of the total simulations (i.e., acceptance rate of 2%) and completed in 22 minutes. All other parameter distributions and results of exploration and stability diagnostics are shown in Supplementary Figure S7. The longer execution time was consistent with the larger number of trajectory-matching time points used in this case (22 compared to 11−13 in earlier models). The influence of the tolerance used to define the feasibility bounds was further evaluated through using larger tolerances of 20% and 30% (SI, Section S11). In this model, runs *M* = 4 was selected as the representative run with the mean energy distance 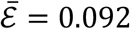.

**Figure 5.**
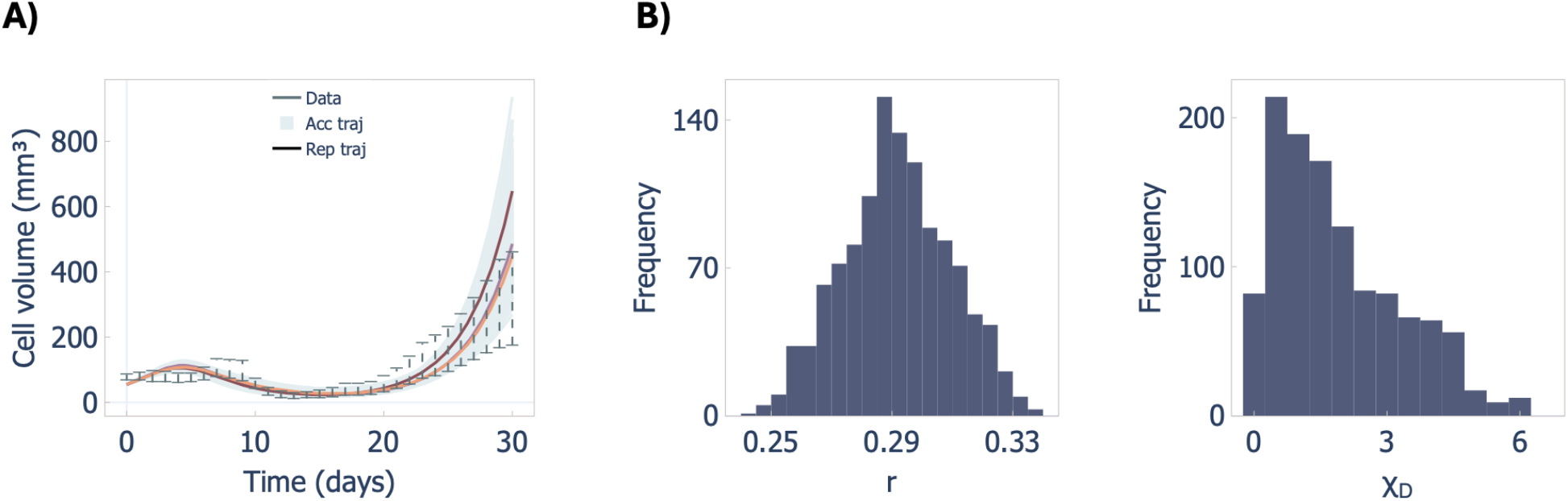
Output of the TM-RWFS algorithm on cancer immunotherapy model. A) Accepted trajectories for the total number of tumor cells based on the TM-RWFS algorithm. B-left) Accepted feasible parameter distribution of tumor growth rate r and B-right) T cell stimulation rate χ_D_ by DCs.

## 4. Discussion and Conclusions

Complex biological systems can exhibit substantial heterogeneity, making it challenging to experimentally or clinically capture the mechanisms driving individual variability. Existing computational methods like ABC-MCMC can help generate such variability but often suffer from high rejection rates^50^, limiting efficiency. Here, we showed that trajectory-matching TM-RWFS generates diverse model trajectories and parameter sets that satisfy data-derived feasibility criteria across biological systems of varying complexity. By replacing the distance-based acceptance step used in ABC variants with a trajectory-matching criterion that retains parameter sets whose outputs fall within specified data bounds, TM-RWFS can reduce rejection under the tested settings. By leveraging best-fit parameter values estimated during model construction for initialization, TM-RWFS begins from a model-informed region of parameter space, which can facilitate exploration but does not guarantee complete coverage of the feasible region. We also assessed the sensitivity of ABC variants to the acceptance threshold *ϵ* and the specification of priors using the Lotka-Volterra model in Case Study 1 and the infection model in Case Study 2A (see Section S10 in the Supplementary Information).

We showed that TM-RWFS accurately and efficiently generates plausible heterogeneous trajectories for biological systems. Across the test cases, TM-RWFS generated accepted feasible parameter populations and trajectories within the specified constraints, with computational performance summarized in (**Table 1** and **Table 2**). This is particularly important as we move towards the integration of TM-RWFS for virtual patient generation, since VPCs must capture the heterogeneity of biological systems to be representative of real clinical populations.

**Table 2.** Summary of TM-RWFS performance for Case studies 2 and 3. For each model, the number of samples (N), number of parallel runs (M), number of data points used for trajectory matching, standard deviation coefficient (C), execution time, number of accepted parameter sets, and acceptance rate (AR) are reported.

| Model | N | M | # Data points | C | Execution time | # Accepted | AR (%) |
| --- | --- | --- | --- | --- | --- | --- | --- |
| Case Study 2A: Influenza infection | $20 \times 10^4$ | 5 | 11 | 0.1 | 13 minutes | 1184 | 0.6 |
| Case Study 2B: Extended influenza infection | $15 \times 10^4$ | 5 | 13 | 0.07 | 10 minutes | 1982 | 0.9 |
| Case Study 3: Cancer immunotherapy | $12 \times 10^4$ | 5 | 22 | 0.1 | 22 minutes | 1173 | 2.2 |

For the immuno-oncology model of Barish et al^40^, TM-RWFS generated tumour trajectories that remained within the specified data bounds at the required time points and showed closer agreement with the experimental measurements during several intervals, including days 1–5, 15–25, and 29–30 (**Figure 6**). These bounds were defined using *mean* ± *SE*, with an additional 10% tolerance applied to *SE*. The resulting accepted feasible parameter distributions (e.g., parameters *c*_*A*_ and *c*_*kill*_, see Figure S7) differed in shape from those reported in the original study, possibly reflecting differences in the parameter regions explored under the TM-RWFS feasibility criteria.

**Figure 6.**
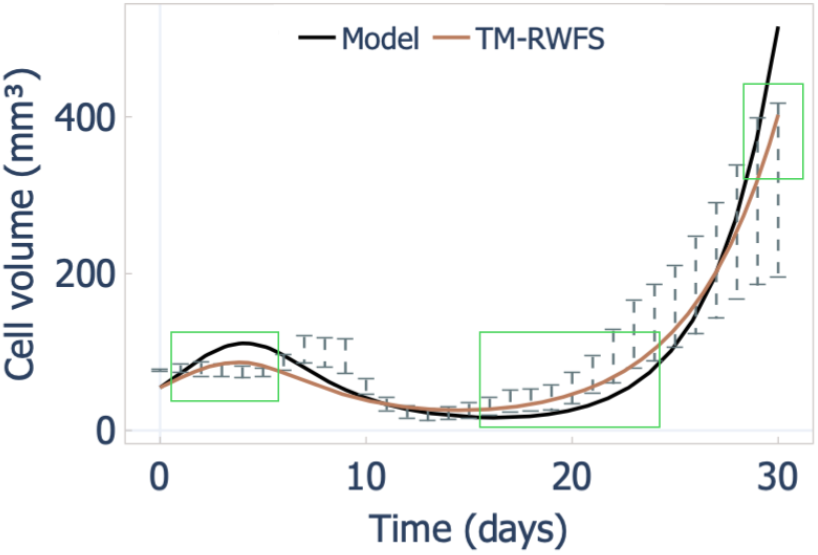
TM-RWFS trajectories showing closer agreement with experimental data at selected time points. TM-RWFS generated tumour trajectories that more closely captured the experimental data during days 1–5, 15–25, and 2S–30, highlighted by the green boxes.

The Barish et al.^40^ immuno-oncology model was hierarchically fit by progressively fixing estimated parameters of a smaller submodel, and subsequently including more equations from which parameters were estimated by matching experimental data. Such a hierarchical approach can influence the resulting parameter estimates because parameters fixed at earlier stages are not jointly re-estimated when additional model components are introduced. Because the credible intervals reported by Barish et al.^40^ were obtained through this hierarchical estimation procedure, we did not interpret their upper and lower limits as independently estimated simultaneous parameter bounds. Instead, these intervals were used to inform the normal proposal distributions, while all parameters were sampled on a linear space (see Supplementary Information for details). Further, the local sensitivity analysis performed by Barish et al.^40^ showed that variations up to 0.7% in *β* result in model fits within 10% of the optimum (SI of Barish et al.^40^). Based on this observation, we evaluated two scenarios, unrestricted and restricted infection rate, with details provided in the SI. We found that constraining the range of β, resulted in greater consistency with the findings of Barish et al.^40^. This difference may reflect the influence of the hierarchical estimation procedure, in which β was estimated at an early stage and subsequently fixed during later fitting steps.

The computational time of TM-RWFS depends on several interacting factors, including the number of measurements used in the trajectory-matching criterion, model complexity, the selected differential equation solver, and the proposal-scaling coefficient *C*. First, as mentioned in the results of Case Study 3 (immuno-oncology), increasing the number of data/time points used during the trajectory matching step increases the execution time of TM-RWFS. Further, because each proposed parameter set must be simulated and assessed against the specified feasibility criteria, the computational cost of solving the model directly affects overall execution time.

Within the tested settings, we also observed that lower values of *C* increased the acceptance rate and reduced the total number of proposals required to obtain the desired number of accepted parameter sets. However, as *C* controls the proposal step size, it therefore affects both acceptance and exploration: a smaller value of *C* produces more local proposals and may increase acceptance while limiting exploration of the feasible parameter space, whereas a larger value allows broader exploration but may result in more frequent rejection. In this study, we used *C* = 0·1 for the influenza infection and cancer immunotherapy models, both with six estimated parameters, and *C* = 0·07 for the extended influenza infection model with 14 estimated parameters, to generate approximately 1000-2000 accepted parameter sets (see Figure S8 for details on the sufficient number of accepted samples).

We used parameter confidence intervals to inform parameter-specific proposal standard deviations, when available. When such intervals were unavailable, constant proposal standard deviations were used instead. The comparison shown in Figure S10 suggests that parameter-specific proposal scales based on available confidence intervals modestly improves sampling performance relative to the tested constant-standard-deviation setting. Nevertheless, the choice of proposal scale can substantially affect sampling behaviour. Therefore, *C* should be treated as a tuning parameter which can influence both acceptance and exploration, for example choosing *C* = 1 will lead to only two accepted parameter sets. *C* = 0·5 gives twenty accepted parameter sets and *C* = 0·25 gives approximately one hundred accepted samples in first infection model for same number of iterations of *N* = 2 × 10^5^. While *C* = 0·1 gives almost 1,000 accepted parameters sets.

In standard Metropolis–Hastings implementations, each proposed sample requires evaluation of the likelihood function, which can be computationally expensive for mechanistic biological models. As a result, although the acceptance rate is typically tuned to lie around 20–30%, each iteration carries a relatively high computational cost^59^. TM-RWFS does not evaluate a likelihood or a Metropolis–Hastings acceptance ratio. Instead, each proposed parameter set is simulated, and the resulting trajectory is assessed directly against the specified feasibility criteria. Consequently, the computational cost of TM-RWFS is determined primarily by model simulation, the number of proposals evaluated, and the trajectory-matching criteria. Because the acceptance mechanisms of Metropolis–Hastings, ABC variants, and TM-RWFS are fundamentally different, their acceptance rates should not be interpreted as directly comparable measures of computational efficiency. In TM-RWFS, a low acceptance rate may arise from restrictive feasibility criteria or large proposal step sizes, whereas a high acceptance rate may result from more local proposals and does not necessarily indicate broader exploration. We therefore considered wall-clock runtime together with the number of simulations required to generate the desired number of accepted feasible parameter sets when evaluating computational performance.

TM-RWFS has two main limitations. First, its performance depends on multiple factors, including the quality of the available clinical or experimental data, the ability of the mechanistic model to capture the data, and the precision of parameter estimation. Second, because TM-RWFS uses local random-walk proposals centred on the most recently accepted parameter values, it may not fully explore all disconnected or widely separated regions of the feasible parameter space. Broader proposal distributions or alternative initialization strategies may improve exploration, but they can also increase rejection rates and computational cost. Indeed, we found that the complex and non-uniform parameter patterns observed for the models studied here were not adequately recovered when candidates were generated using uniform distributions with fixed bounds. This reflects differences in how parameter space exploration and does not imply that uniform distributions are intrinsically inappropriate for feasibility sampling, under the correct conditions. Future improvements to TM-RWFS could therefore focus on strategies that enhance exploration, particularly in scenarios with limited data or poorly constrained parameter estimates, thereby broadening its potential application to virtual patient cohort generation (see Section S8 in the Supplementary Information).

Although TM-RWFS was initialized using previously estimated best-fit parameter values in the primary analyses, initialization at the best-fit parameter set is not strictly required. We additionally tested initialization with half of the parameters randomly set away from their best-fit values. Under this test, TM-RWFS still generated accepted feasible parameter sets and trajectories (Figure S11). These results suggest that TM-RWFS can tolerate moderate perturbations of the initial parameter values, although the starting point may affect exploration of the feasible region. Therefore, sensitivity to initialization should be assessed, particularly for models with complex or potentially disconnected feasible parameter spaces.

In summary, TM-RWFS generated accepted feasible parameter populations and heterogeneous model trajectories across mechanistic models of varying complexity, with execution times on the order of minutes for the cases examined in this study. Although direct computational comparisons with other virtual patient cohort generation methods depend on model structure, implementation, hardware, and acceptance criteria, these results demonstrate that TM-RWFS can provide a computationally practical approach for generating feasible parameter sets and trajectories. This may be particularly useful when large numbers of heterogeneous model realizations are required for model-based analyses. Overall, TM-RWFS provides a trajectory-based feasibility sampling framework for representing biologically plausible variability in mechanistic mathematical models, with applications for virtual patient cohort generation and virtual clinical trial analyses.

## ACKNOWLEDGMENTS

This work was funded by the Fondation CHUSJ and a Vanier Canada Graduate Scholarship (SHC), NIH NIAID R01 AI170115 (AMS and MC) and, in part, thanks to the Canada Research Chair in Computational Immunology with funding from the Canada Research Chairs Program (MC).

## AUTHOR CONTRIBUTIONS

Conceptualized the research: FB, JJAW, SHC, CJS, AMS, MC

Designed the algorithm: FB and MC

Performed the research: FB

Wrote the manuscript: FB and MC

Edited the manuscript: FB, JJAW, SHC, CJS, AMS, MC

## CODE AVAILABILITY

The code to run Trajectory Matching-Random Walk Feasibility Sampling (TM-RWFS) and to generate the figures in this paper is available at: https://github.com/Craig-Lab/TM-RWFS.

