## Supplementary Information for "Trajectory matching-random walk feasibility sampling for simulating heterogeneous dynamics in mechanistic models"

### S1. Comparison of ABC and TM-RWFS for the Lotka-Volterra model

| Parameter | Description | Best fit value (Orig) | ABC | TM-RWFS | |
|---|---|---|---|---|---|
| $a$ | prey growth rate | 1.0 | [0.99, 1.12] | [0.99,1.1] | [0.88, 1.16] |
| $b$ | predator reproduction rate | 1.0 | [0.85, 1.13] | [0.88, 1.07] | [0.8, 1.2] |
| $Initial\ conditions\colon\ x(0) = 1, y(0) = 0.5$ | | | | | |

***Table S1. Estimations for Case Study 1: Lotka-Volterra model.*** *Parameters of the Lotka–Volterra model[1] with best-fit values used in TM-RWFS and the initial conditions. Parameter ranges were estimated using ABC (for $\epsilon \leq 4.3$) and TM-RWFS: left column, $\epsilon \leq 4.3$; right column, $\epsilon \leq 5.5$.*

### S2. Parameter estimates for Case Study 2A: Influenza infection

| Parameter | Description | Unit | Best fit | 95% CI | Range (Curr) |
|---|---|---|---|---|---|
| $\beta$ | Virus infectivity | $\mathrm{TCID}_{50}^{-1}\ \mathrm{d}^{-1}$ | -3.61 | [-4.3, -1.1] | [-4.77, -1.97] |
| $p$ | Virus production | $\mathrm{TCID}_{50}^{-1}\ \mathrm{cell}^{-1}\mathrm{d}^{-1}$ | 0.2 | [-0.08, 2.09] | [-0.18, 3.22] |
| $c$ | Virus clearance | $\mathrm{d}^{-1}$ | 1.1 | [0.8, 2.9] | [0.75, 4.24] |
| $k$ | Eclipse phase | $\mathrm{d}^{-1}$ | 4 | [4.0, 6.0] | [4.0, 5.99] |
| $\delta_d$ | Infected cells clearance | $\mathrm{cell}^{-1}\mathrm{d}^{-1}$ | 6.2 | [6.14, 6.23] | [6.14, 6.29] |
| $K_\delta$ | Half Saturation constant | cells | 5.05 | [2.08, 5.2] | [2.73, 5.52] |
| $Initial\ conditions\colon\ T(0) = 10^7, I_1(0) = 75, I_2(0) = 0, V(0) = 0$ | | | | | |

***Table S2. Estimations for Case Study 2A: Influenza infection.*** *Parameters along with their best-fit values used in TM-RWFS for the density-dependent viral kinetics model. The 95% confidence intervals from the original study were used to define the standard deviations of the normal distributions and the initial conditions[2]. Parameter ranges were estimated using TM-RWFS. All parameters are reported in logarithmic scale, except for $k$.*

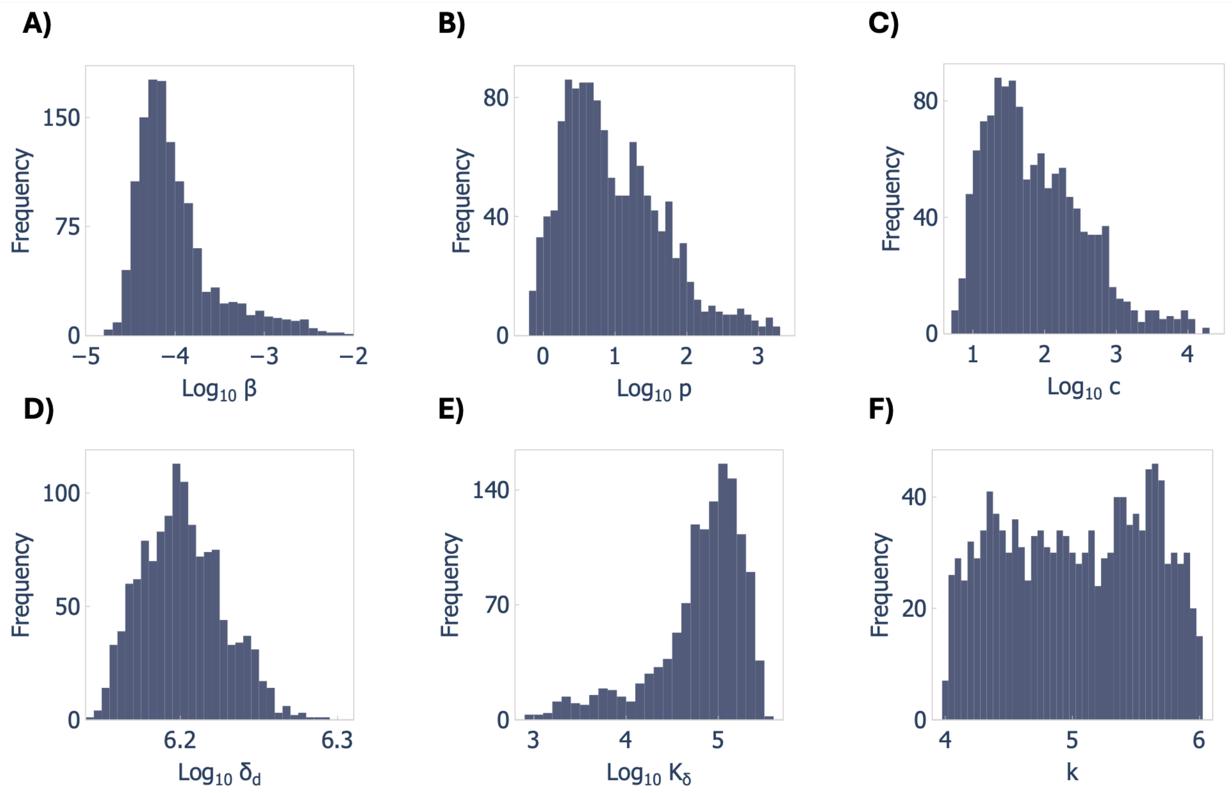


***Figure S1. Accepted feasible parameter distribution of Case Study 2A: Influenza infection.*** *A) Virus infection rate* $\beta$*, B) virus production rate* $p$*, C) virus clearance rate* $c$*, D) infected cells clearance rate* $\delta_d$ *, E) half saturation constant* $K_\delta$ *, F) Eclipse phase rate* $k$.

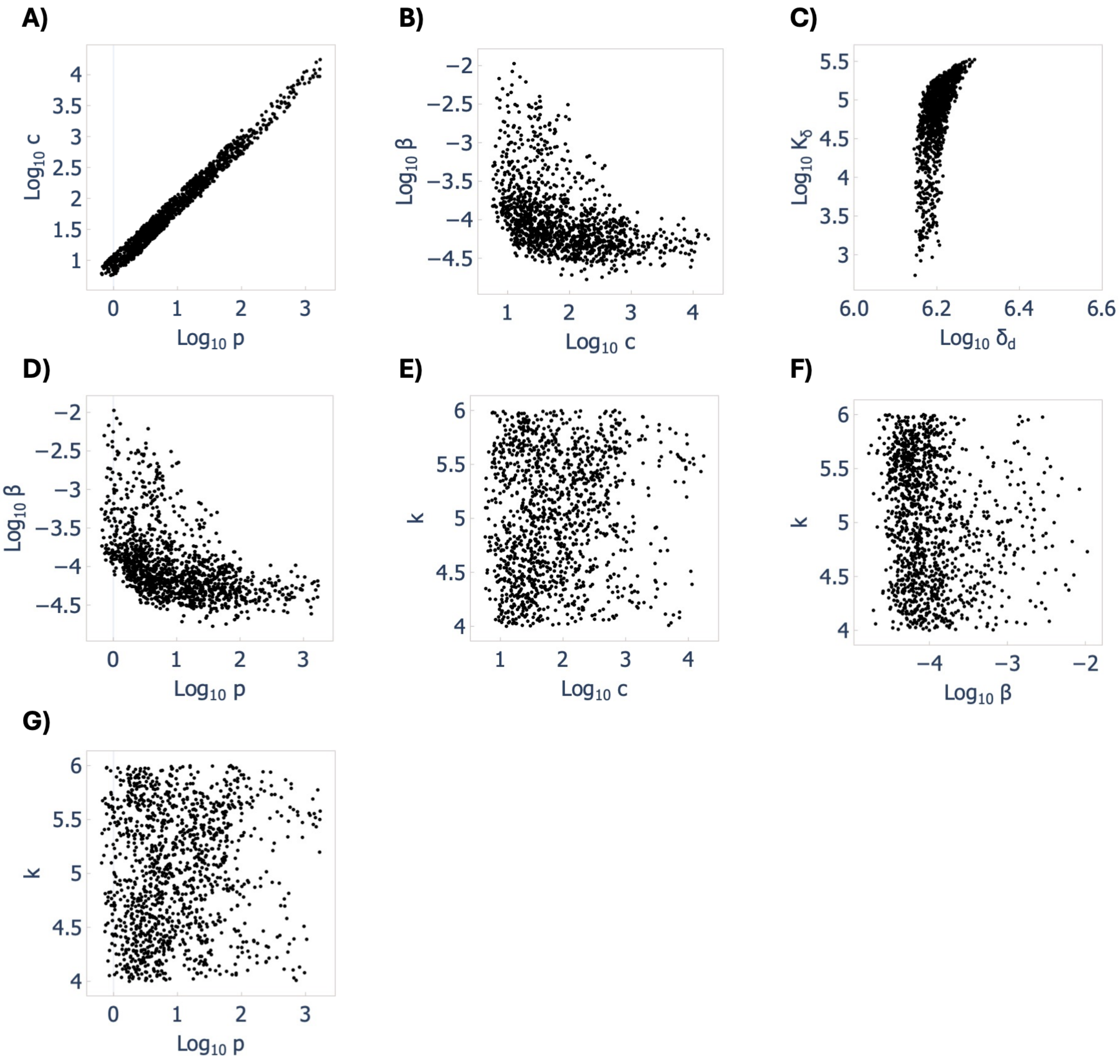


***Figure S2. Scatter plots of parameter estimate of Case Study 2A: Influenza infection.*** *Scatter plots of parameter pairs which show positive correlations between the virus production rate, $p$ and the virus clearance rate, $c$ (A), consistent with the original study*[2].

### S3. Parameter estimates for Case Study 2B: Influenza infection

| Parameter | Description | Unit | Best fit | 95% CI | Range (Curr) |
|---|---|---|---|---|---|
| $\beta$ | Virus infectivity | $\mathrm{TCID}_{50}^{-1}\ \mathrm{d}^{-1}$ | -4.2 | [-5.3, -4.0] | [-5.55, -4.03] |
| $p$ | Virus production | $\mathrm{TCID}_{50}^{-1}\ \mathrm{cell}^{-1}\mathrm{d}^{-1}$ | 0.0 | [0.23, 2.04] | [-0.22, 2.46] |
| $c$ | Virus clearance | $\mathrm{d}^{-1}$ | 0.97 | [0.74, 2.97] | [0.73, 2.99] |
| $k$ | Eclipse phase transition | $\mathrm{d}^{-1}$ | 4.0 | [4.0, 6.0] | [4.0, 5.99] |
| $\delta$ | Infected cells clearance | $\mathrm{d}^{-1}$ | 0.24 | [0.1, 0.66] | [0.07, 0.69] |

| | | | | | |
|---|---|---|---|---|---|
| $\delta_E$ | Infected cells clearance by $CD8_E$ | $\mathrm{d}^{-1}$ | 1.9 | [0.33, 2.0] | [0.15, 1.99] |
| $K_{\delta_E}$ | Half-saturation constant | cells | 2.6 | [2.0, 5.4] | [1.0, 5.60] |
| $\xi$ | $CD8_E$ infiltration | $CD8_E^2$ $\mathrm{cell}^{-1}\mathrm{d}^{-1}$ | 4.4 | [2.1, 4.9] | [2.04, 5.91] |
| $K_E$ | Half-saturation constant | $CD8_E$ | 5.9 | [3.0, 6.0] | [3.0, 6.9] |
| $\eta$ | $CD8_E$ expansion | $\mathrm{cell}^{-1}\mathrm{d}^{-1}$ | -6.6 | [-7.8, -6.2] | [-7.37, -6.2] |
| $\tau_E$ | Delay in $CD8_E$ expansion | d | 3.6 | [2.1, 5.9] | [2.0, 5.10] |
| $d_E$ | $CD8_E$ clearance | $\mathrm{d}^{-1}$ | 1.0 | [0.05, 2.0] | [0.07, 1.99] |
| $\zeta$ | $CD8_M$ generation | $CD8_M$ $CD8_E^{-1}$ $\mathrm{d}^{-1}$ | 0.22 | [0.01, 0.94] | [0.01, 0.96] |
| $\tau_M$ | Delay in $CD8_M$ generation | d | 3.5 | [3.0, 4.0] | [3.0, 3.9] |
| $Initial\ conditions$: $T(0) = 10^7, I_1(0) = 75, I_2(0) = 0, V(0) = 0, E(0) = 0, E_{M1}(0) = 0$ | | | | | |

***Table S3. Parameter estimates for Case Study 2B: Influenza infection.*** *Parameters along with their best-fit values used in TM-RWFS for the extended model of influenza infection including $CD8^+$ $T$ cell dynamics. The 95% confidence intervals from the original study were used to define the standard deviations of the normal distributions and the initial conditions. The parameter ranges were estimated using TM-RWFS. All parameters are reported in logarithmic scale, except for $k$, $\delta, \delta_E, \tau_E, d_E, \tau_M$* and *$\zeta$. In TM-RWFS, parameter constraints were applied while running TM-RWFS according to Myers et al[3].*

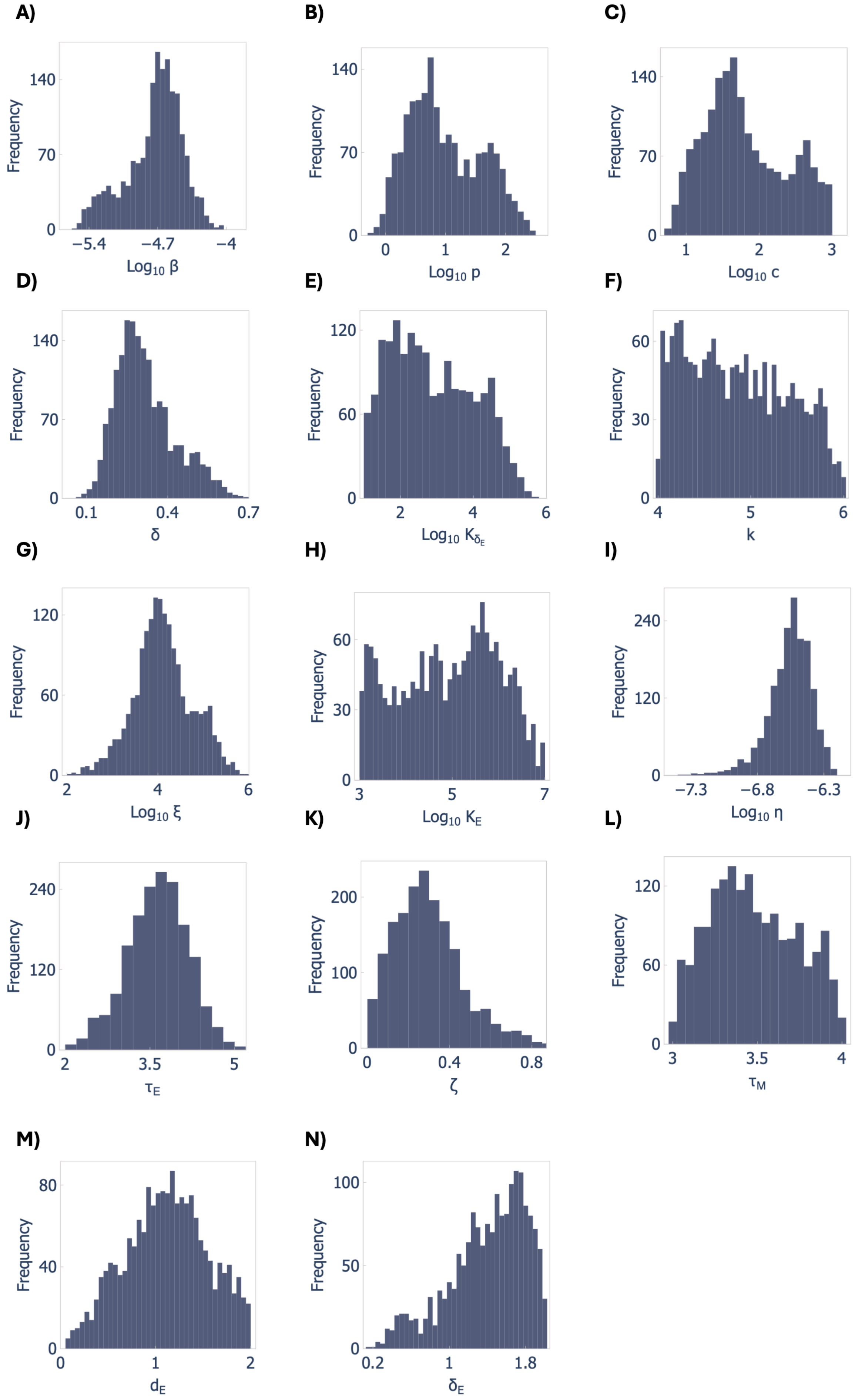
A)
Frequency
140
70
0
−5.4
−4.7
−4
$\text{Log}_{10}\ \beta$
B)
Frequency
140
70
0
0
1
2
$\text{Log}_{10}\ p$
C)
Frequency
140
70
0
1
2
3
$\text{Log}_{10}\ c$
D)
Frequency
140
70
0
0.1
0.4
0.7
$\delta$
E)
Frequency
120
60
0
2
4
6
$\text{Log}_{10}\ K_{\delta_E}$
F)
Frequency
60
30
0
4
5
6
k
G)
Frequency
120
60
0
2
4
6
$\text{Log}_{10}\ \xi$
H)
Frequency
60
30
0
3
5
7
$\text{Log}_{10}\ K_E$
I)
Frequency
240
120
0
−7.3
−6.8
−6.3
$\text{Log}_{10}\ \eta$
J)
Frequency
240
120
0
2
3.5
5
$\tau_E$
K)
Frequency
200
100
0
0
0.4
0.8
$\zeta$
L)
Frequency
120
60
0
3
3.5
4
$\tau_M$
M)
Frequency
80
40
0
0
1
2
$d_E$
N)
Frequency
100
50
0
0.2
1
1.8
$\delta_E$

***Figure S3. Accepted feasible parameter distribution of Case Study 2B: Influenza infection.*** *A) Infection rate $\beta$, B) virus production rate $p$, C) virus clearance rate $c$, D) Infected cells clearance rate $\delta$, E) half-saturation constant $K_{\delta_E}$, F) eclipse phase transition rate $k$, G) $CD8_E$ infiltration rate $\xi$, H) half-saturation constant $K_E$, I) $CD8_E$ expansion rate $\eta$, J) Delay in $CD8_E$ expansion $\tau_E$, K) $CD8_M$ generation rate $\zeta$, L) delay in $CD8_M$ generation $\tau_M$, M) $CD8_E$ clearance rate $d_E$, N) Infected cells clearance rate by $CD8_E$, $\delta_E$.*

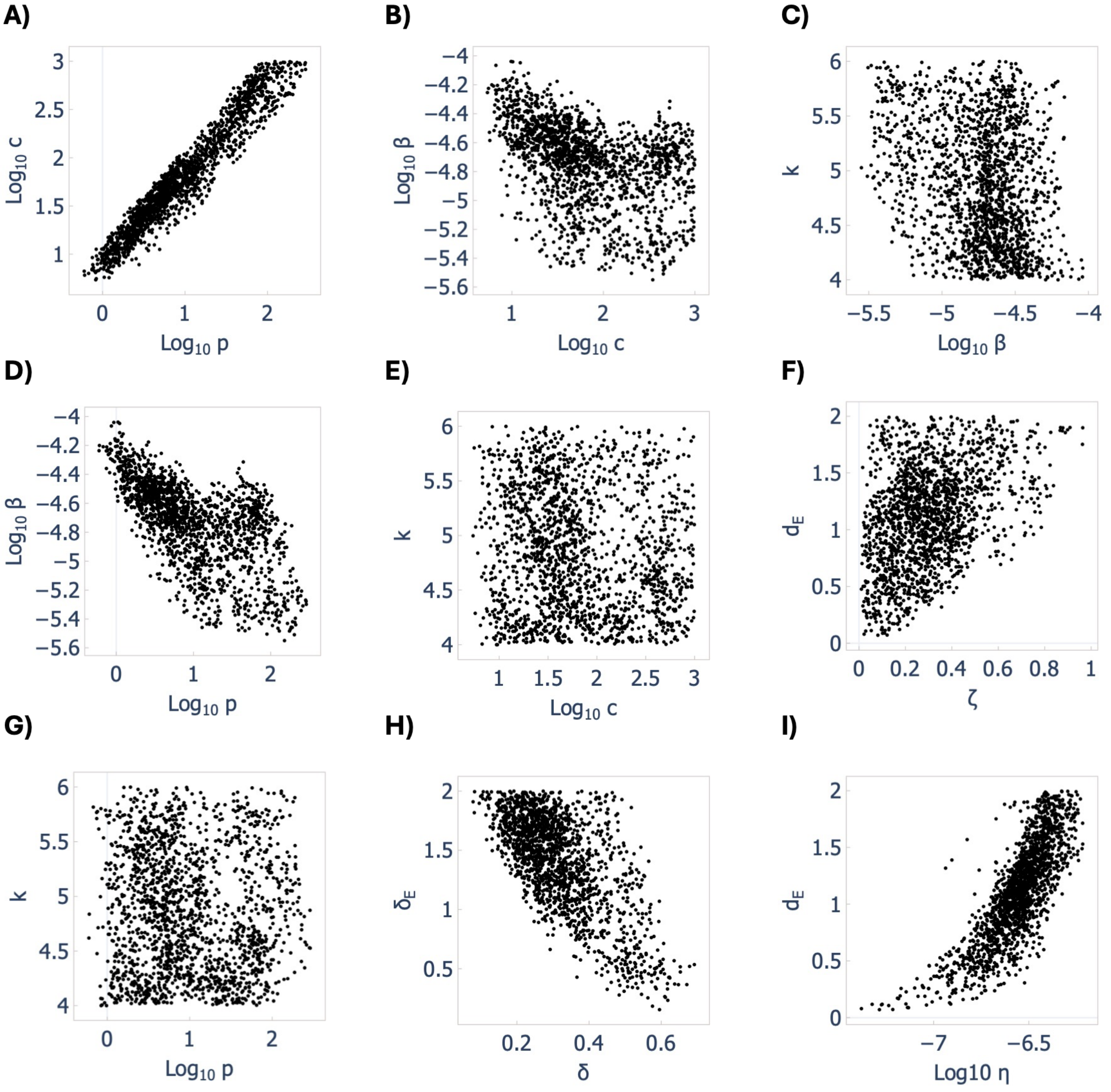


***Figure S4. Scatter plots of parameter estimate of Case Study 2B: Influenza infection.*** *Scatter plots of parameter pairs which show positive correlations between the virus production rate,* $p$ *and the virus clearance rate,* $c$*, (A) and slightly negative correlation between the virus clearance rate,* $c$*, and the infection rate,* $\beta$*, (B), between the virus production rate,* $p$*, and the infection rate,* $\beta$*, (D), between the infected cells clearance rate by* $CD8_E$*,* $\delta_E$*, and the infected cells clearance rate,* $\delta$*, (H) and correlation between* $CD8_E$ *clearance rate,* $d_{E,}$ *and* $CD8_M$ *generation rate,* $\zeta$*, (I), consistent with the original study[3].*

### S4. Case Study 3: Cancer immunotherapy

To define the sampling space necessary to run TM-RWFS (see Section 3.1, $Eq.(10)$ and $Eq.(11)$ in the Main Text), we tested both logarithmic and linear spaces for all six model parameters (i.e., $r,\ \beta, \chi_D, c_A, c_T, and\ c_{kill}$) and found that sampling all parameters from their linear values yielded more reliable results based on energy distance metric (see Section 3.2 in the Main Text). Moreover, we evaluated two scenarios when applying TM-RWFS: (1) no restriction on the accepted values of the infection rate ($\beta$), and (2) restricting $\beta$ to lie within

($\beta^* \pm 0.15\,\beta^*$), where $\beta^*$ denotes the best-fit value estimated in Barish et al.[4] Note that the 15% margin was selected arbitrarily to limit excessive variability while preserving sufficient flexibility for exploring the parameter space and generating diverse trajectories.

| Parameter | Description | Unit | Best fit | 95% Credible interval | Range (Curr) |
|---|---|---|---|---|---|
| **Fixed parameters** | | | | | |
| $\alpha$ | Viral production | $virions$ | 3000 | … | … |
| $\delta_I$ | Infected lysis rate | $\mathrm{d}^{-1}$ | 1.0 | … | … |
| $\delta_V$ | Viral decay rate | $\mathrm{d}^{-1}$ | 2.3 | … | … |
| $\delta_T$ | T-cell decay rate | $\mathrm{d}^{-1}$ | 0.35 | … | … |
| $\delta_A$ | Naïve T-cell decay rate | $\mathrm{d}^{-1}$ | 0.35 | … | … |
| $k_0$ | Base T-cell killing rate | $\mathrm{d}^{-1}$ | 2.0 | … | … |
| $\chi_A$ | T-cell differentiation rate | $\mathrm{d}^{-1}$ | 1.0 | … | … |
| $\delta_D$ | Dendritic cell death rate | $\mathrm{d}^{-1}$ | 0.35 | … | … |
| **Estimated parameters** | | | | | |
| $r$ | Net tumor growth rate | $\mathrm{d}^{-1}$ | 0.32 | [0.29, 0.36] | [0.24, 0.33] |
| $\beta$ | Infection rate | $\mathrm{d}^{-1}$ | $1\times10^{-3}$ | [$0.9\times10^{-3}$, $1\times10^{-3}$] | [$0.95\times10^{-3}$, $1.04\times10^{-3}$] |
| $c_A$ | Naïve T-cell recruitment rate | $\mathrm{d}^{-1}$ | $5\times10^{-4}$ | [$2\times10^{-4}$, 1.66] | [$5\times10^{-4}$, 0.91] |
| $c_T$ | T-cell recruitment rate | $\mathrm{d}^{-1}$ | 1.2 | [$8\times10^{-3}$, 4.15] | [$8\times10^{-3}$, 5.7] |
| $c_{kill}$ | Enhanced T-cell cytotoxicity | $\mathrm{cell}^{-1}\mathrm{d}^{-1}$ | $5.1\times10^{-7}$ | [$1\times10^{-4}$, 1.1] | [$3\times10^{-3}$, 2.32] |
| $\chi_D$ | Rate of T-cell stimulation by DCs | $\mathrm{d}^{-1}$ | 5.5 | [4.02, 8.47] | [$4\times10^{-3}$, 6.2] |

| $Initial\ condition: U(0) = 60, I(0) = 0, V(0) = 0, T(0) = 0, A(0) = 0, D(0) = 0$ |
|---|

***Table S4. Estimations for Case Study 3: Cancer immunotherapy**. Parameters along with their best-fit values used in TM-RWFS for the* model of tumour growth after combination immunotherapy. *The 95% credible intervals from the original study were used to define the standard deviations of the normal distributions and the initial conditions[4]. Parameter ranges were estimated using TM-RWFS. All parameters are reported in linear scale.*

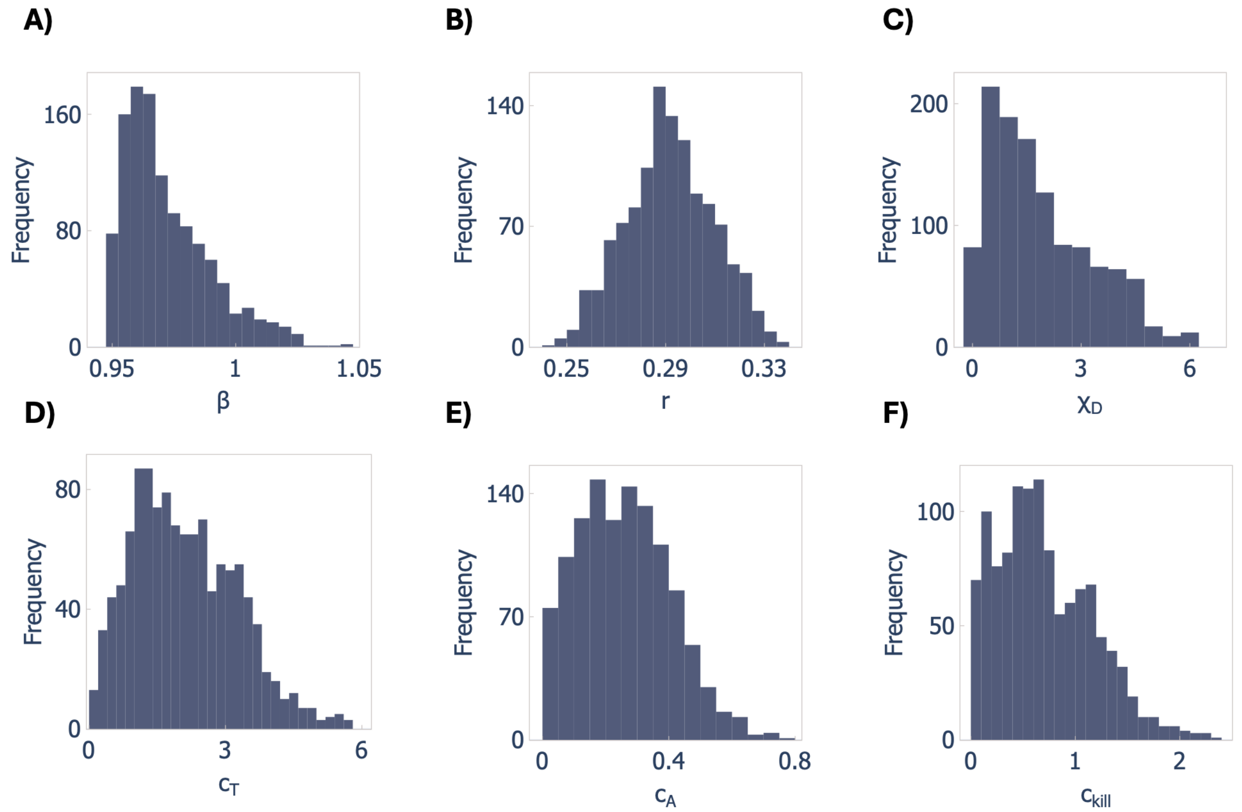


***Figure S5. Accepted feasible parameter distribution of Case Study 3: Cancer immunotherapy.** A) Virus replication rate $\beta$, B) tumour growth rate $r$, C) T-cell stimulation by DCs rate $\chi_D$, D) T-cell recruitment rate $c_T$, E) naïve T-cell recruitment rate $c_A$, F) enhanced T-cell cytotoxicity rate $c_{kill}$ .*

**S5. Effect of increasing sample size on stability of accepted parameter distribution**

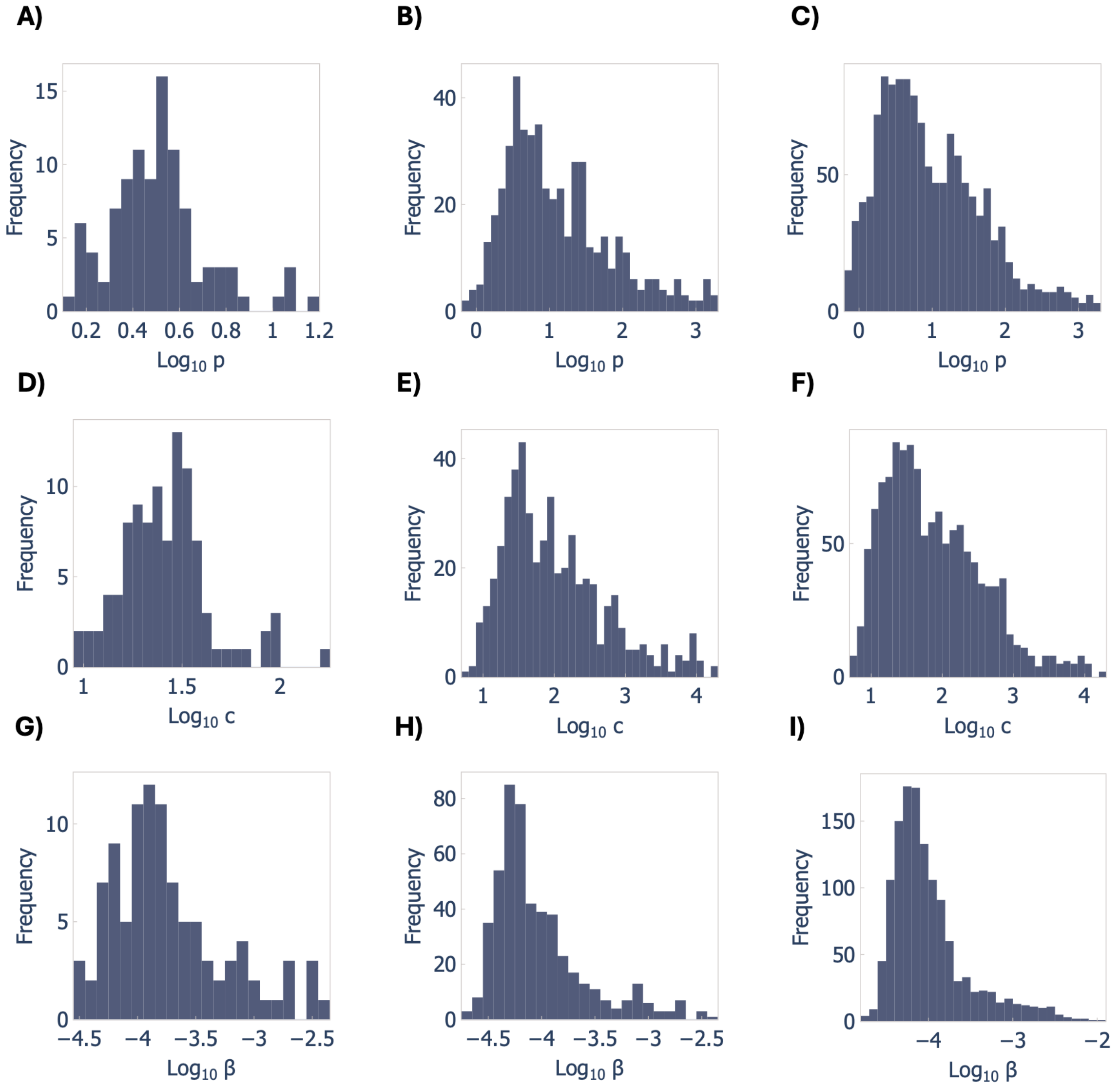


***Figure S6. Effect of accepted-sample size ($N_A$) on stability (infection model, case study 2A).*** *A-C: histograms of virus production rate, p for* $N_A$ = 100, 500, 1000. *D-F: virus clearance rate, c; G-I: infection rate,* $\beta$.

To investigate how accepted distribution stability depends on accepted-sample size ($N_A$) in TM-RWFS, we examined three sizes of accepted samples_100, 500, and 1000_in the density-dependent viral-kinetics model for parameters $p$, $c$, and $\beta$. **Figure S6** shows how the posterior distributions of $p$, $c$, and $\beta$ evolve as the number of accepted samples increases from 100 to 1000. With only $N_A = 100$, the distributions appear noisy and unstable, lacking smooth structure. As $N_A$ increases to 500 and then 1000, the histograms become smoother and more stable, indicating improved convergence and greater reliability of the inferred distributions.

This highlights the importance of continuing sampling until a sufficient number of accepted trajectories is reached to ensure robust inference.

**S6. Importance of time point selection in TM-RWFS**

Through application to the Lotka-Volterra model (see Section 4.1 in the Main Text), we showed that when extrema are well characterized and model dynamics are oscillatory, generating a large number of acceptable trajectories with TM-RWFS can be achieved on the order of seconds versus standard ABC methods that may require several hours. However, careful attention should be paid to the choice of time points selected for the trajectory matching step so that they correspond to the dynamical features of interest. Indeed, significant changes to a system's behavior (i.e., change in slope direction, biphasic rates etc.) are particularly critical for capturing informative dynamics and improving inference efficiency. To assess how the choice of observation timepoints influences both the inferred trajectories and the efficiency of TM-RWFS, we varied the time-point selection in the infection model (Case Study 2A) and examined the resulting trajectories. The dataset has 11 observation times; the main text uses all 11 within TM-RWFS. Here, we also evaluate alternative schedules, including a) only the first few time points; b) only the last few time points; c) only the middle time points; d) omission of the middle time points; e) dispersed/irregularly spaced time points; and f) other feasible combinations. The combinations of selected time points along with TM-RWFS performance and resulting trajectories are summarized in **Table S5** and **Figure S7**. Except for Case D, the predicted trajectories exhibited poor convergence, with samples arising from different regions of the parameter space, resulting in diverse and inconsistent dynamics. Underlining the importance of identifying critical time points, though Case D did not include all observations but nonetheless covered the initial, final, and transitional time points where the model dynamics undergo important changes (e.g., the viral load peaks around time point 2 and then declines slowly; this slow decay transitions to a faster decay around time points 7–8). This suggests that omitting less informative time points can still yield satisfactory results while reducing computation time. Although some of the generated trajectories may not be biologically meaningful, their importance ultimately depends on the specific purpose for which the trajectories are generated. For example, if we are interested in patients whose viral load dynamics peak above the maximum value (~6), we can refer to the parameter values corresponding to such a trajectory in Case B of **Figure S7**.

| Case | N | # Data points | Execution time | Accepted | Convergence |
|---|---|---|---|---|---|
| A | $5\times10^3$ | 0,1,2,3 | 13 s | 1563 | No |
| B | $2\times10^4$ | 7,8,9,10 | 60 s | 1079 | No |
| C | $5\times10^3$ | 0,1,8,9 | 15 s | 1741 | No |
| D | $2\times10^5$ | 0,1,2,7,8,9,10 | 4.6 minutes | 935 | Yes |
| E | $1\times10^4$ | 0,3,6,9 | 44 s | 2177 | No |
| F | $2\times10^4$ | 0,2,4,6,8,10 | 67 s | 1678 | No |
| G | $5\times10^3$ | 0,1,9,10 | 18 s | 1636 | No |
| H | $2\times10^4$ | 3,4,5,6 | 53 s | 4787 | No |

***Table S5. Summary of TM-RWFS performance under different time-point selections in the infection model (Case Study 2A).*** *For each case, the number of samples ($N$), data points used for trajectory matching, execution time, number of accepted parameter sets, and convergence results are reported.*

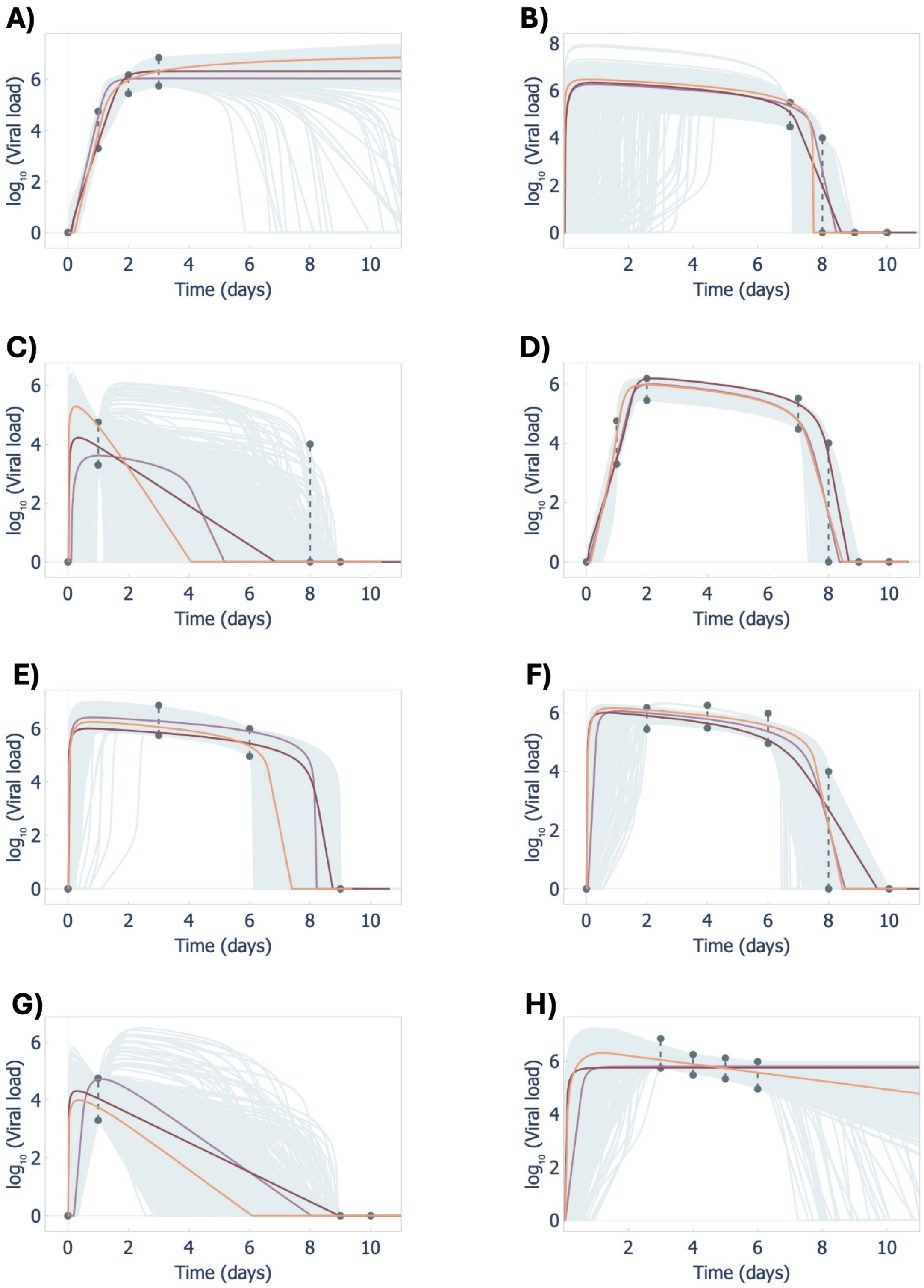


***Figure S7. Accepted trajectories under different time-point selections in the infection model (Case Study 2A).*** *Virus dynamics trajectories when timepoints A) [0,1,2,3], B) [7,8,9,10], C) [0,1,8,9], D) [ 0,1,2,7,8,9,10], E) [0,3,6,9], F) [ 0,2,4,6,8,10], G) [ 0,1,9,10] and H) [3,4,5,6] are selected for TM-RWFS.*

### S7. TM-RWFS results without using confidence intervals

Throughout the paper, we use confidence intervals (CIs) or credible intervals (CrIs) to determine the standard deviations of Normal distributions (Section 3.1, Eq. (5)), where $\sigma_i$ is derived from the difference between the upper and lower bounds which are determined using CIs or CrIs. Here, we consider the case in which no CI/CrI is available, and only by best-fit parameter values are reported. We previously tested this scenario in the Lotka–Volterra model by assigning a fixed standard deviation to each parameter's normal distribution. Here we repeat the experiment on a more complex system to evaluate TM-RWFS performance. Specifically, in the infection model (Case Study 2A) we set $\sigma_i = 0.1$ for all six parameters ($i = 1, \ldots, 6$). We showed that the generated trajectories and distributions are satisfactory (**Figure S8**), but the exploration of parameter space is narrower than in the main case (**Table S7**), where 95% CIs were used to define the standard deviations, resulting in different standard deviations for different parameters. However, the algorithm completed in ~3 minutes using fewer iterations and producing 1,800 accepted samples, which was more efficient than the main case (**Table S6**). Thus, there is a trade-off between parameter space exploration and execution time; When heterogeneity matters (which is generally the case for virtual patient cohorts), we can use parameter-specific SDs derived from CIs/CrIs; the runtime will be longer and the number of accepted samples smaller, but the parameter space is explored more thoroughly. When heterogeneity is less critical and the goal is simply to find alternative parameter sets that fit the data beyond the best-fit vector, we can use equal SDs across parameters for faster execution at the cost of narrower exploration.

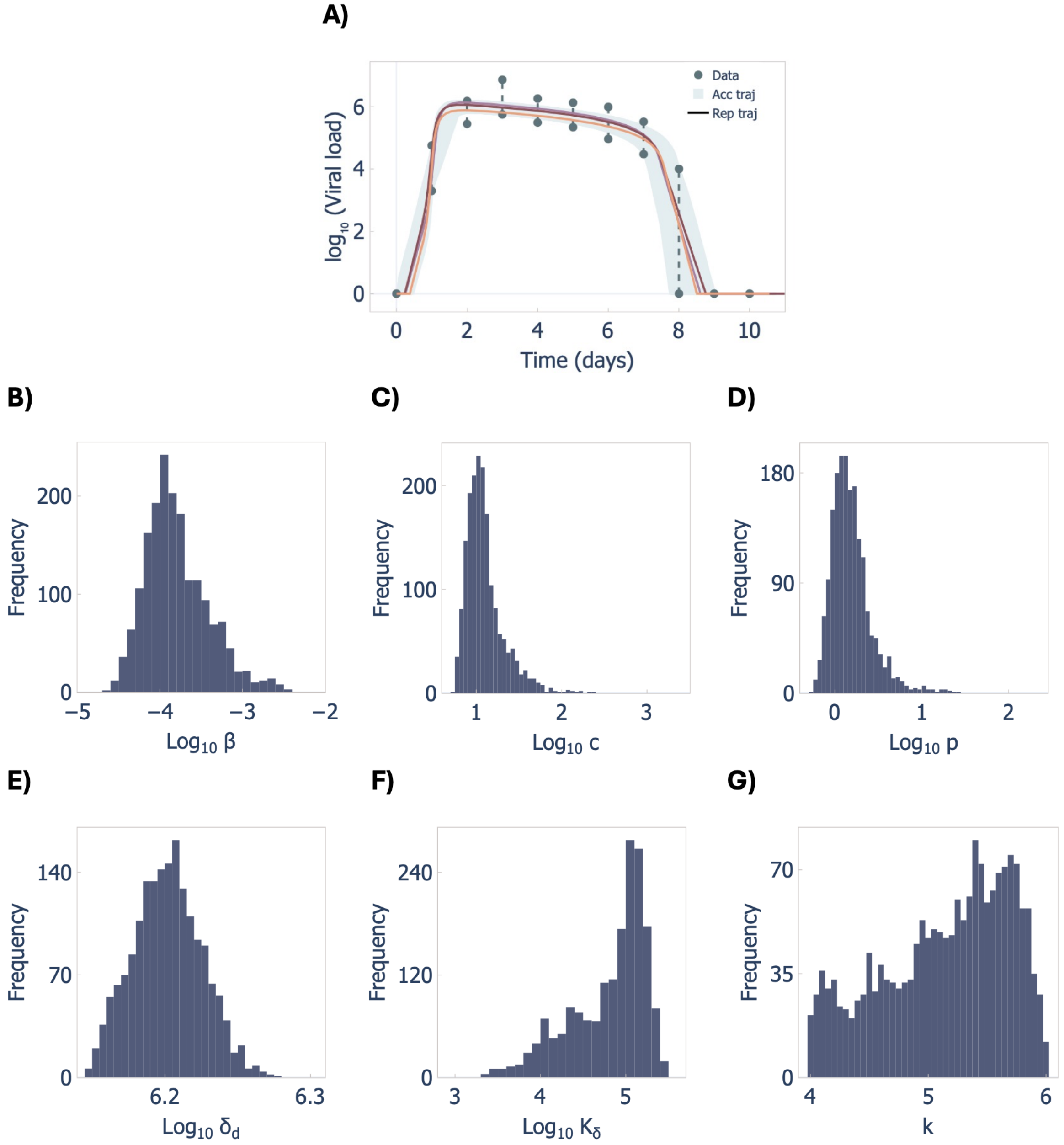


***Figure S8. Accepted trajectories and parameters distributions in TM-RWFS without using CIs in the infection model (Case Study 2A).*** *A) Virus-dynamics trajectories, B-G) parameters distributions. Note that for parameters $\beta$,$p$,$c$ and $K_\delta$, the full range is not fully covered by the accepted samples.*

| N | # Data points | Execution time | Accepted | Convergence |
|---|---|---|---|---|
| $5\times10^4$ | 11 | 3 minutes | 1800 | Yes |

***Table S6. Summary of TM-RWFS performance without using CIs in the infection model (Case Study 2A).*** *The number of samples (N) and data points used for trajectory matching, execution time, number of accepted parameter sets, and convergence result.*

| Parameter | Range (using 95% CIs)<br>$\sigma_\beta = \mathbf{0.32}$ , $\sigma_p = \mathbf{0.13}$ , $\sigma_c = \mathbf{0.33}$,<br>$\sigma_k = \mathbf{0.2}$, $\sigma_{\delta_d} = \mathbf{0.1}$ , $\sigma_{K_\delta} = \mathbf{0.32}$ | Range (fixed SD)<br>$\sigma = \mathbf{0.1}$ |
|---|---|---|
| $\beta$ | [-4.87, -2.09] | [-4.6, -2,4] |
| $p$ | [-0.25, 2.55] | [-0.26, 1.4] |
| $c$ | [0.7, 3.32] | [0.7, 2.3] |
| $k$ | [4.0, 5.99] | [4, 5.9] |
| $\delta_d$ | [6.14, 6.29] | [6.14, 6.27] |
| $K_\delta$ | [2.81, 5.54] | [3.31, 5.45] |

***Table S7. Comparing the ranges of parameters in infection model (Case Study 2A) for fixed and varying standard deviations.*** *The range for parameters $\beta$, $p$, $c$ and $K_\delta$ are smaller for fixed standard deviation compared to varying standard deviations.*

### S8. TM-RWFS performance with altered prior parameter estimates

To explore the TM-RWFS performance when best fit values are not provided for all the parameters of a mathematical model, we examined the infection model (Case Study 2A) under the scenario in which the best-fit values of half of the parameters were deliberately changed. Specifically, we arbitrarily selected the parameters $\{k, \beta, K_\delta\}$ and perturbed the parameters relative to the best-fit values: $k$ $+25\%$ $(4 \rightarrow 5)$, $K_\delta$ $-40\%$ $(5.05 \rightarrow 3$ , logarithmic scale) and $b$ decreased in magnitude by $\sim 17\%$ $(-3.62 \rightarrow -3.00$ , logarithmic scale). These values were then used as the means of the normal distributions (see Section 3.1, $Eq.(4)$ in the Main Text) to initialize TM-RWFS. Despite not starting at previously estimated best fit values, TM-RWFS still generated parameter distributions and trajectories consistent with those reported by Smith et al[2]. This demonstrates that TM-RWFS remains effective even when some parameter estimates are sloppy, as long as they lie within biologically meaningful ranges (**Figure S9**).

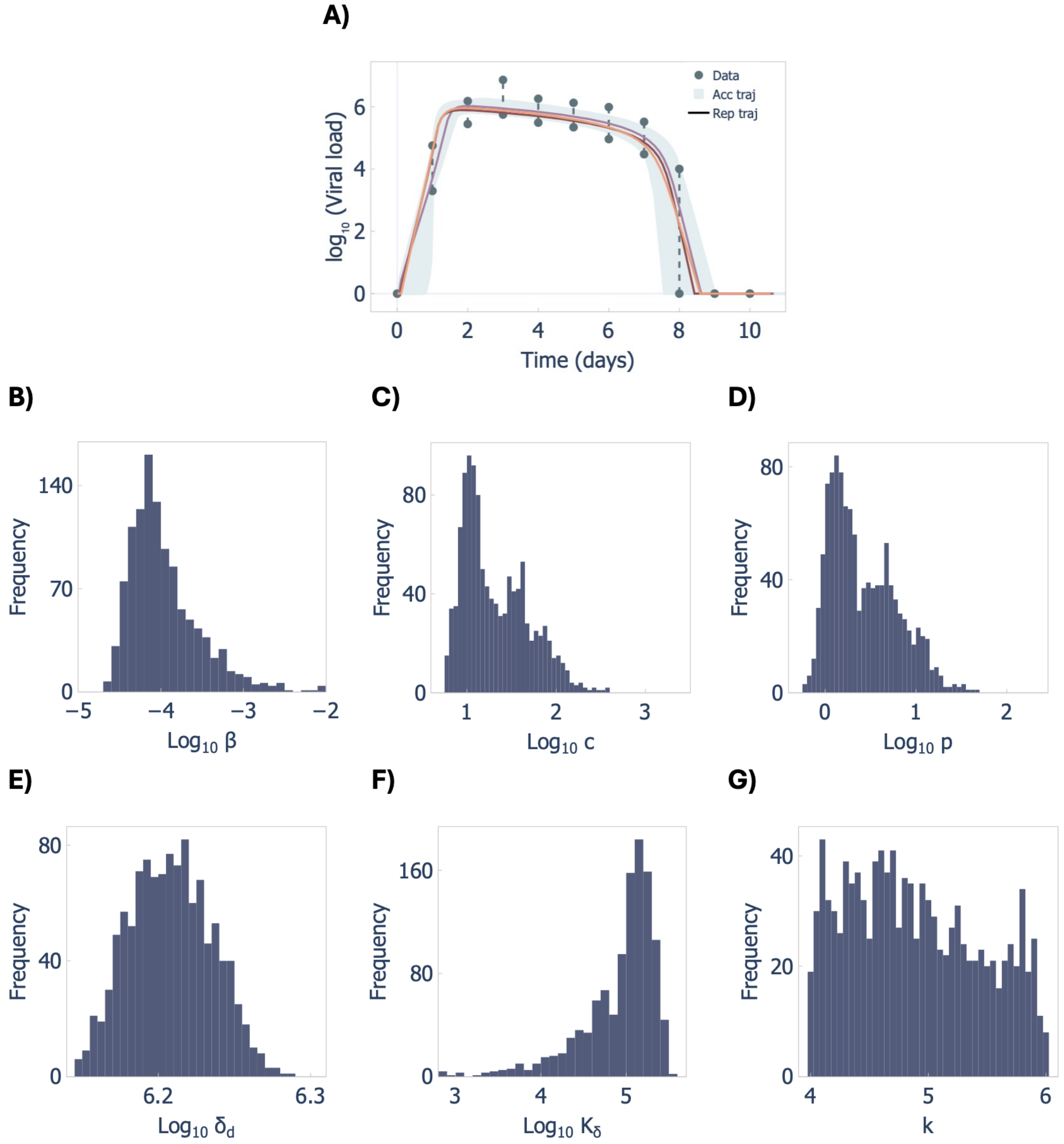


***Figure S9. Accepted trajectories and parameters distributions in TM-RWFS with perturbed parameters in the infection model (Case Study 2A).*** *A) Virus-dynamics trajectories, B-G) parameters distributions where parameters $k, \beta, K_\delta$ are perturbed.*

### S9. Generating synthetic data from population-level data and model prediction

If individual level data or standard errors are not explicitly reported at each measured time point, some strategies can be employed to construct uncertainty bounds for data:

**A. Scaling the observed data.** A fixed percentage (e.g., $\pm 20\%$) is applied directly to the observed data values. The data bounds are then calculated (see Section 2.3, $Eq. (7.a)$ and $Eq. (7.a)$ in the Main Text) as:

$$x_i^{upper} = D_i + 0.2D_i \quad and \quad x_i^{lower} = D_i - 0.2D_i.$$

**B. Mirroring around the model predicted best fit values.** Construct symmetric uncertainty bounds by reflecting each data point above/below the predicted trajectory. Specifically, let $x_i^{pred}$ be the model prediction and $D_i$ the data observed at time $t_i$. Then, the lower bound can be defined as:

$$x_i^{lower} = x_i^{pred} - \left(D_i - x_i^{pred}\right) = 2\, x_i^{pred} - D_i,$$

while the upper bound remains the observed data point $D_i$, or vice versa. This ensures that the model prediction lies within the constructed bounds (**Figure S12.B**).

As shown in **Figure S10**, the method in Case A does not guarantee that the model prediction will fall within the constructed range. In the illustrated case, the model trajectory lies within the bounds at only three points (1.1, 7.5, and 9.6). Thus, requiring a simulated trajectory during TM-RWFS to be within the presented bounds at all eight time points may not be feasible. To compare the results of these two strategies (**Figure S10**), we first reproduced the results of ABC on LV model reported in Toni et al.[1], then we implemented TM-RWFS on this model using the methods in Cases A and B.

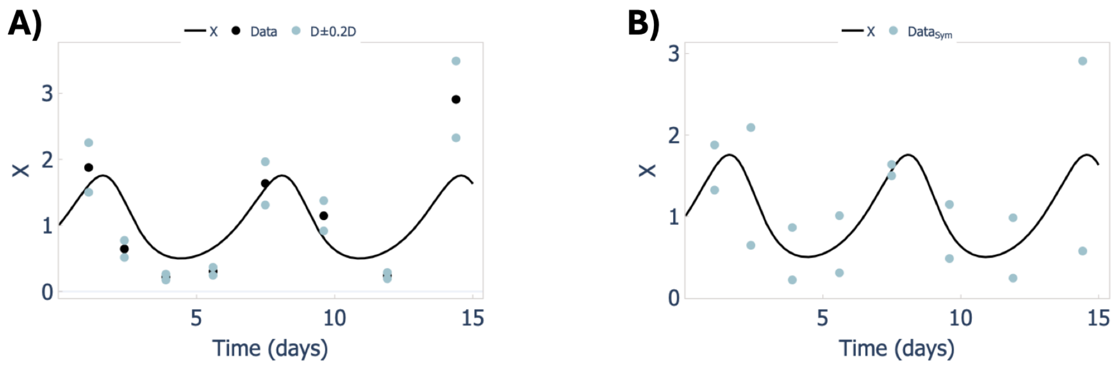


***Figure S10. Different approaches to generate data uncertainty using Lotka-Volterra model.*** *A) Scaling the observed data. B) Mirroring around the model predicted best fit values.*

To fairly compare both cases, we selected same time points for each strategy. In Case A, parameter $a$ does not fully explore the parameter space and the distribution of parameter $b$ does not overlap with the distributions obtained from ABC (**Figure S11, B**). This is to be expected, since none of these parameter sets satisfied the condition $\epsilon \leq 4.3$ (**Table S8**). In Case B, there is partial overlap between the distributions, while the non-overlapping portion may be because only four time points were used in TM-RWFS (**Table S8**). If all eight time points had been included, the degree of overlap would likely have increased (**Figure S11, D**).

| Case | N | Data type | # Data points | Execution time | All accepted | Accepted $\epsilon \leq 4.3$ |
|---|---|---|---|---|---|---|
| A | $5\times10^4$ | D$\pm$0.2D | 1,5,6,8 | 42 s | 1531 | 0 |
| B | $15\times10^4$ | $Data_{sym}$ | 1,5,6,8 | 12 s | 2256 | 1098 |

***Table S8. Summary of TM-RWFS performance on LV model under two data-uncertainty scenarios.*** *A) Scaling the observed data. B) Mirroring around the model predicted best fit values.*

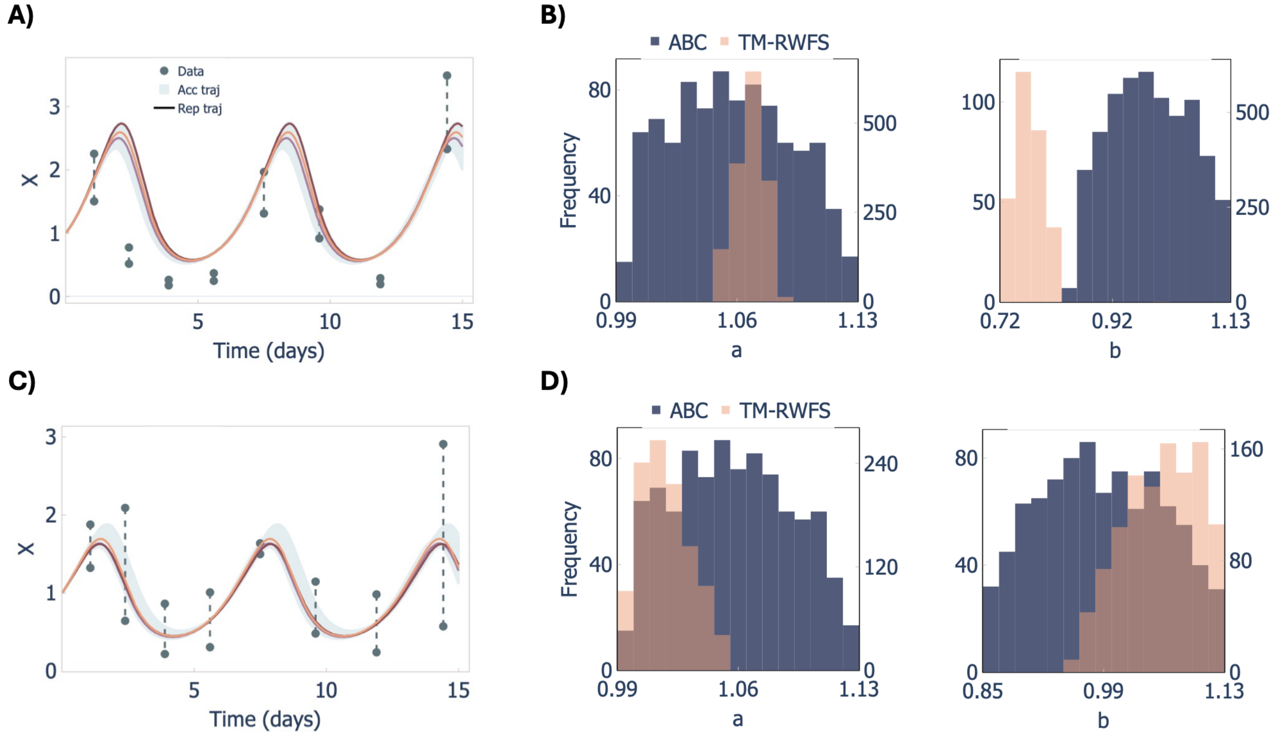


***Figure S11. Accepted trajectories and parameter distributions in the LV model*** *under two data-uncertainty scenarios.* A-B) *D ± 0.2D, where none of the trajectories satisfied* $\epsilon \leq 4.3$. C-D) *$Data_{sym}$, showing 1,098 accepted samples with* $\epsilon \leq 4.3$.

### S10. Implementing Approximate Bayesian Computation Sequential Monte Carlo (ABC-SMC) in the Lotka Volterra and influenza infection models

ABC-SMC is a potent and popular variant of ABC that employs an efficient sampling scheme by maintaining a population of parameter “particles” drawn from the prior[1,5-7]. At each stage and with decreasing tolerance, particles are weighted based on the resulting distance between simulated model predictions and observed data, and higher-weight particles are resampled and perturbed to form the next population, which progressively approximating the posterior[1,7-9]. pyABC was developed by Klinger et al.[5,10] and is a Python implementation of ABC-SMC with adaptive thresholds, configurable distance functions, and parallel execution. We compared pyABC's performance with TM-RWFS in the infection model (Case Study 2) and LV model (Case Study 1). However, because of differences in the implementation of the two algorithms, we first adjusted the tolerance threshold $\epsilon$ and population size in pyABC to match those used in TM-RWFS. From the trajectories accepted using TM-RWFS, we obtained $\epsilon_{min} = 0.8$, $\epsilon_{max} = 4.4$, and $\bar{\epsilon} = 2.5$. Thus, we considered two pyABC configurations in which the distance function for the virus-dynamics trajectories, defined as the sum of absolute errors (SAE) in the Ordinary Differential Equations section of the pyABC package, was required to be less than or equal to the final (minimum) tolerance, with $\epsilon_{final}$ set to 2.5 and 4.4.

Since in TM-RWFS, we produce about 1000 accepted samples, we configured pyABC with a population size of 1000 for comparability. As the parameter $k$ must lie in the biologically plausible range $[4,6]$, we tested two pyABC setups:

1. Informative prior: $k \sim Uniform(4,6)$, enforcing the constraint directly.
2. Non-informative prior with post-filtering: increase the population size from 1000 to 4000, use $k \sim Uniform(0,10)$, then discard particles with $k \notin [4,6]$ ($\approx 75\%$ removed).

At $\epsilon = 4.4$ (the maximum acceptance threshold in TM-RWFS), pyABC had a faster execution (4 minutes versus 13 minutes for TM-RWFS) using an informative prior for $k$ to enforce biologically plausible bounds. However, this came at the cost of poorly resolved posterior distributions, particularly for $\beta$ **(Figure S12A)** and, to a lesser extent, $\delta_d$. Using a stricter threshold ($\epsilon = 2.5$), pyABC remained faster (5 minutes versus 13 minutes for TM-RWFS) under the same informative prior on $k$ and provided accurate posterior distribution for $\beta$. In contrast, when a noninformative prior was used, the pyABC runtime increased to approximately 5 minutes at $\epsilon = 4.4$. Applying an additional post hoc filtering step to remove parameter sets with $k \notin [4,6]$ reduced biologically implausible solutions but again led to less accurate posterior distributions for $\beta$ **(Figure S12C)**. At $\epsilon = 2.5$ with a noninformative prior for $k$, the runtime further increased to 16 minutes; however, after filtering out parameter sets with $k \notin [4,6]$, all parameters retained accurate posterior distributions. Overall, these results indicate that computational efficiency in pyABC depend strongly on tolerance range and how prior knowledge about $k$ is incorporated either directly through an informative prior or indirectly via post-processing filters to exclude biologically implausible parameter values for $k$. The distributions of parameter $\beta$ across different scenarios of infection model are shown in **Fig. S12.**

| | $\epsilon = 4.4$ | $\epsilon = 2.5$ |
|---|---|---|
| Informative prior for parameter k | A) 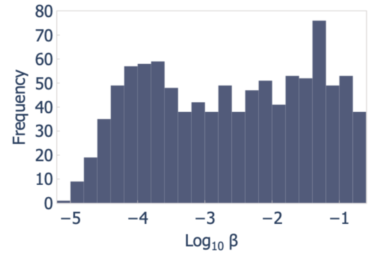 | B) 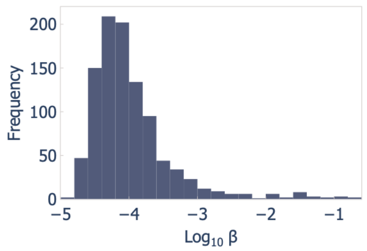 |
| Noninformative prior for parameter k | C) 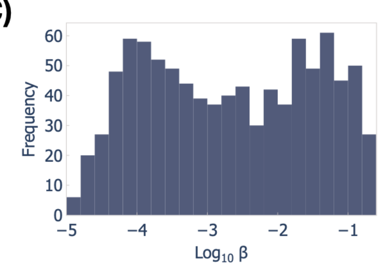 | D) 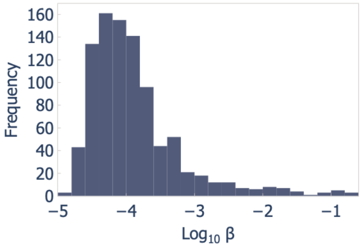 |



***Figure S12. Histograms of the viral infection rate (β).*** *A(B) Informative prior for eclipse phase rate k and* $\epsilon = 4.4$ *(*$\epsilon = 4.4$*), C(D) Informative prior for eclipse phase rate k and resampling after pyABC to keep parameter sets with* $k \in [4{,}6]$ *for* $\epsilon = 4.4$.

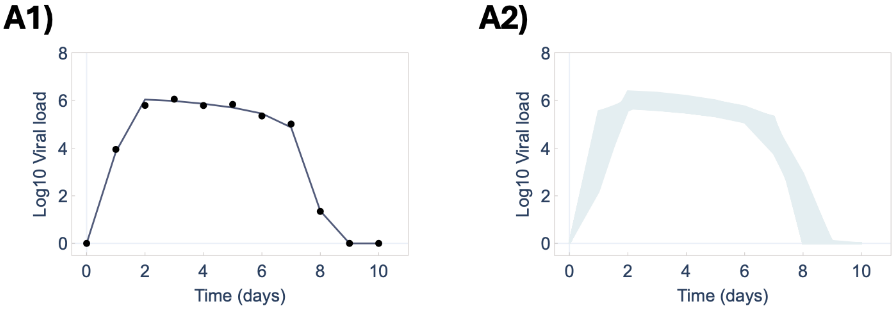

***Figure S13. pyABC results in accurate model predictions for infection model.*** *A1) Viral load dynamics prediction using best fit parameter values (solid black), data (black dots) with* $\epsilon \approx 0.9$. *A2) Accepted trajectories of viral load using pyABC.*

We then applied pyABC to the LV model, where the execution time was longer than that of TM-RWFS (75 seconds versus 8 seconds). In addition, the inferred range of parameter $b$ differed from that reported in Toni et al.[1] for ABC-SMC: the range in pyABC was substantially broader, extending up to approximately $4$ (**Fig.S14, B4**), whereas Toni et al.[1] report values below $1.4$. It is important to note that part of this discrepancy arises from differences in the definition of the distance function and the amount of data used in pyABC framework versus the implantation Toni et al. used for ABC-SMC. In Toni et al.[1], the distance was defined using the SSE across both components of the model, whereas in the pyABC implementation it is defined using SAE based on only one component of the system. This difference in the distance metric and data leads to parameter distributions that are not directly comparable between the two approaches. Resulting distributions for parameters of LV model using pyABC are shown in **Fig. S14, B3-B4 Fig. S14.B2**. As can be seen from the corresponding trajectories shown in **Fig. S14.B2**, this mismatch results in predictions that are less aligned with the data.

One way to improve both the inferred parameter distributions and the resulting trajectories is to incorporate more data into the pyABC. For example, the distributions and trajectories reported in Toni et al.[1] are more accurate and biologically meaningful because ABC-SMC was applied using data from both components of the system. A reason why in general pyABC performs better in the infection model compared to the LV model, despite the increased complexity of the infection model, is the robustness of the model itself. Indeed, the previously performed parameterization of the infection model shows the model to be a very close fit to the data[2] (with $\epsilon \approx 0.9$, **Fig.S13, A1**), facilitating the generation of parameter sets that closely reproduce the observed dynamics. Furthermore, in pyABC, the model is evaluated only at the specific time points where measurements are available. This approach works well as long as the model aligns closely with the observed data. Comparing the predicted trajectories of the LV model (**Fig.S14, A2** with $\epsilon \approx 4.3$) to the infection model (**Fig.S13, A2**) resulting from ABC-SMC demonstrates that the trajectories in the latter case are much better aligned with the data, whereas there is a substantial mismatch between the model and the data for the LV model. Under these conditions, evaluating the model only at specific measurement time points, as done in pyABC, can degrade the algorithm's performance and may contribute to the reduced accuracy of the inferred distributions (**Fig.S14, B3 and B4**).

To increase accuracy, the ODE solver in pyABC can be modified so that the model is evaluated on a dense time grid per simulation rather than only at the measurement times (e.g., the 8 time points by default for LV model). The simulated trajectories can then be interpolated to the measurement times before comparison with the data. This modification increased the computational cost per simulation, resulting in a longer total runtime (approximately 130 seconds), but improved the acceptance rate and the accuracy of parameters distributions. **Fig. S14** illustrates the results obtained using dense time evaluation (**A1-A4**) compared to those based on evaluation only at the measured data points (**B1-B4**). The results show that both the trajectories and the inferred parameter distributions are improved for dense time evaluation and more closely match those reported in Toni et al.[1]

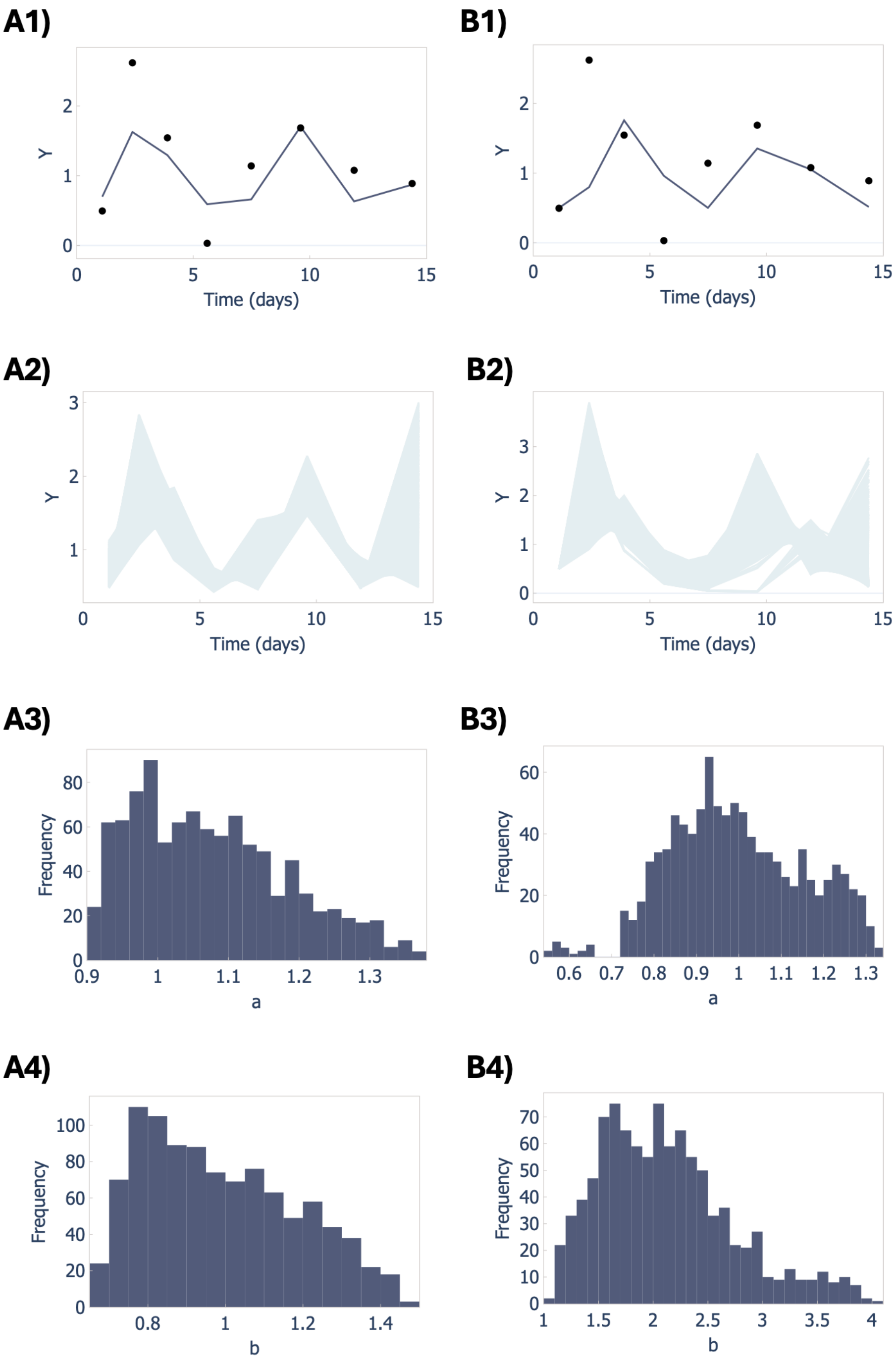


***Figure S14. pyABC results for the LV model.** Predator dynamics prediction (solid black), data (black dots) with evaluating the model at dense time grid (A1) or at specific data points (B1), with corresponding accepted trajectories (A2) and (B2) respectively and the parameters distributions of A3-A4 and B3-B4 respectively.*

### S11. The influence of different choices of tolerances in Case study 3

We further evaluated the outputs of the algorithm by altering the tolerance added to the standard error in Case study 3 to 20% and 30%. Results show increased number of accepted samples and more heterogeneous trajectories for tumor volume compared to the 10% tolerance. Choosing a specific tolerance depends on the application and the amount of variability one wants to have in generated trajectories. **Figure S15** shows the tumor volume trajectories and accepted parameters distributions for 20% and 30% tolerances with 2748 and 3494 accepted parameters respectively. Due to larger flexibility in accepted trajectories, the accepted range of the parameter's values will also increase accordingly.

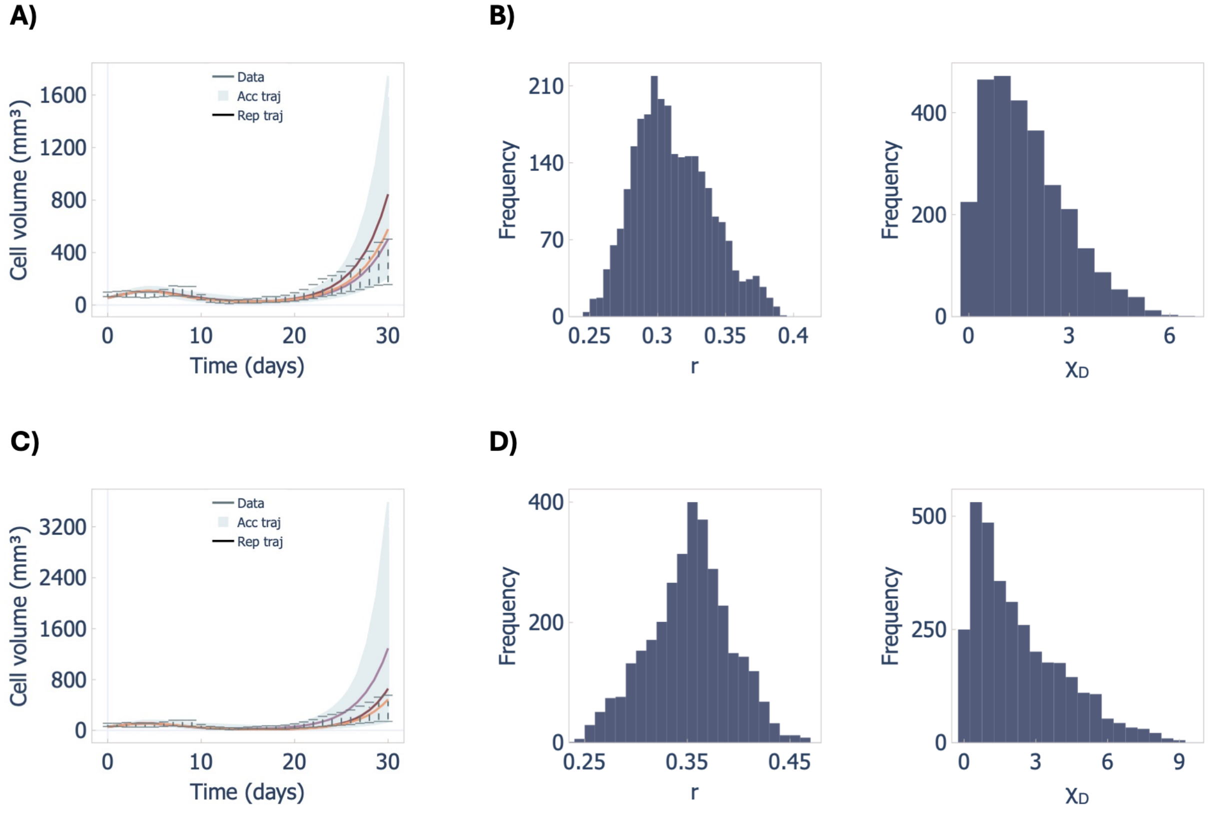


***Figure S15. The influence of different choices of tolerances to define the feasibility bounds in Case study 3****. A-B) Accepted tumor volume trajectories and parameters distributions for 20% tolerance and C-D) for 30% tolerance.*

#### S12. Wasserstein distance

Agreement across repeated TM-RWFS runs can also be evaluated using Wasserstein distance to assess the marginal distributional agreement for each parameter. Pairwise Wasserstein distances are given by:

$$W_1(F,G) = \int_{-\infty}^{\infty} |F(z) - G(z)|\, dz$$

where $z$ denotes the parameter value used for comparison, and $F(z)$ and $G(z)$ are the corresponding cumulative distribution functions from the two TM-RWFS runs. For each run, the mean Wasserstein-1 distance, $\overline{W}_1$, was calculated relative to the remaining runs. In Case study 2A, runs $M = 4$ and $M = 5$ produced highly comparable results. For $M = 4$, $\bar{W}_1 = 0.404$ and the corresponding mean energy distance was $\bar{\mathcal{E}} = 0.683$, whereas for $M = 5$, $\bar{W}_1 = 0.423$ and $\bar{\mathcal{E}} = 0.631$. In Case Study 2B, runs $M = 2$ and $M = 5$ produced highly comparable results. For $M = 2$, $\bar{W}_1 = 0.228$ and $\bar{\mathcal{E}} = 0.199$ , whereas for $M = 5$, $\bar{W}_1 = 0.224$ and $\bar{\mathcal{E}} = 0.213$. In Case Study 3, both energy and Wasserstein distances identified $M = 4$ as the representative run.
